# Ageing-related defects in macrophage function are driven by *MYC* and *USF1* transcriptional programmes

**DOI:** 10.1101/2023.09.20.558643

**Authors:** Charlotte E. Moss, Simon Johnston, Martha Clements, Veryan Codd, Stephen Hamby, Alison H. Goodall, Sumeet Deshmukh, Ian Sudbery, Daniel Coca, Heather L. Wilson, Endre Kiss-Toth

## Abstract

Macrophages are central innate immune cells whose function declines with age. The molecular mechanisms underlying age-related immunity changes remain poorly understood, particularly in human macrophages. We report a substantial reduction in phagocytosis, migration and chemotaxis in human monocyte-derived macrophages (MDMs) from older (>50 years) compared with younger (18-30 years) donors, alongside downregulation of transcription factors *MYC* and *USF1* with age. In MDMs from young donors, knockdown of *MYC* or *USF1* decreased phagocytosis and chemotaxis and altered expression of genes associated with these functions, as well as adhesion and extracellular matrix remodelling. A concordant dysregulation of *MYC* and *USF1* target genes was also seen in MDMs from older donors. Furthermore, older age and loss of either *MYC* or *USF1* in MDMs led to an increased cell size, altered morphology and reduced actin content. Together, these results define *MYC* and *USF1* as key drivers of MDM age-related functional decline and identify downstream targets to improve macrophage function in ageing.

## Introduction

Macrophages are a critical component of the cell-mediated immune system and act as both regulatory and effector cells in healthy and disease contexts. Macrophages have diverse functions at the interface of the innate and adaptive immune systems, including chemotaxis and migration through tissues, phagocytosis of pathogens and other foreign material, clearance of dead and dying cells, antigen presentation, and cytokine production (1). Macrophages exist as many sub-types, depending on tissue environment, differentiation and activation. The typing of macrophages into pro-inflammatory M1 and anti-inflammatory M2 has been invaluable in understanding their biological function *in vitro*. However, much work has also gone into understanding tissue-resident macrophage function, showing that the M1 and M2 classification is much too simplistic for *in vivo* studies. Macrophages exhibit versatility in different settings and nomenclature should focus on stimulating agents or macrophage function (2).

Macrophages have been demonstrated to have a key role in inflammaging, the chronic, sterile inflammatory phenotype that has been intensely studied in ageing research (3), and have been implicated in functional immune decline with age and are involved in many age-related diseases such as atherosclerosis, diabetes, fibrosis, immunocompromise, autoimmunity and cancer (4). It is thought that macrophages become chronically activated with age, with their function more closely resembling an inflammatory macrophage phenotype, including increased inflammatory and stress responses (5). However, how the molecular changes that determine the chronic inflammatory phenotype translate to a decline in function is unclear, a critical gap limiting the development of new immunotherapeutics for ageing. A further barrier to new therapeutics is our current reliance on mouse models, with a lack of studies in humans (6). Here, we sought to measure ageing-related changes in macrophage migration and phagocytosis in a human cohort as a phenotypic marker to identify the molecular changes underlying macrophage ageing.

Age-related changes in phagocytosis and chemotaxis have been reported in several contexts in the literature, mainly using murine models, but with differing outcomes. Aged mouse peritoneal macrophages show reduced phagocytosis with diverse phagocytic targets including apoptotic neutrophils (7), fluorescent particles (8) and myelin fragments (9). In contrast, phagocytosis of zymosan particles by peritoneal macrophages was found to be similar between young and old animals (10), while an earlier study found that phagocytosis activated by adjuvants occurred equally across different age groups (11). Studies of brain resident macrophages (microglia) revealed that uptake of myelin is reduced with age (9,12), as was uptake of lipid debris (13). Similarly, alveolar macrophages consistently show a reduction in phagocytic activity with age, including lower phagocytosis of apoptotic neutrophils (14) and *E. coli* (15). However, aged mouse bone marrow-derived macrophages (BMDM) showed no defect in uptake of zymosan particles (10), fluorescent particles (8), latex particles (16), or *E. coli* (17). In contrast, decreased phagocytosis with age was seen for uptake of apoptotic Jurkat cells (18), myelin fragments (9) and apoptotic N2A cell debris or *E. coli* bioparticles (19). To date, no publications have directly assessed the effect of age on phagocytosis in human monocyte-derived, primary macrophages (MDMs).

Mouse peritoneal macrophages and BMDMs showed improved chemotaxis with age, including towards chemoattractants FMLP (20) and CCL2 (21) and in an *in vivo* study measuring the number of antigen presenting cells migrating to the lymph nodes from the site of injection (22). Contrastingly, aged BMDMs had slower migration towards an implanted polypropylene mesh surface (17). Therefore the effects of aging were dependent on the environment, which determines the macrophage phenotype, as well as by the nature of stimulus.

Few publications have studied the effects of aging on migration in humans. Aged human GM-CSF-derived MDMs showed preferential migration towards conditioned media from immune senescent cells (23). The early macrophage response in human macrophages to pro-inflammatory stimuli such as lipopolysaccharide (LPS), a bacterial cell wall component, is well understood and utilised in both *in vivo* and *in vitro* models. LPS stimulation has also been well studied in ageing contexts, where macrophages from older animals have tissue-specific alterations in pro-inflammatory responsiveness (24). Although LPS induces a predominantly M1 phenotype, more varied phenotypes may exist *in vivo* and other phenotypic changes and stimuli have been less studied.

Transcription factor (TF) analysis has been key in interpreting macrophage biology and function, but so far has only been used in a small number of studies that focussed on inflammaging. In aged murine macrophages, loss of diurnal rhythmicity of transcription factor KLF4 has been attributed to loss of diurnal rhythmicity of phagocytosis with age (25). Transcription factor regulons were used to predict that KLF4 and IRF5 were decreased with age in tissue macrophage populations. This was associated with a trend to a switch from an M1-like to an M2-like macrophage phenotype, known to occur in age-related tumour growth promotion (26). Older alveolar macrophages have been found to have a senescent-like phenotype with reduced proliferation, insensitivity to growth factor stimulation, and increased p16 and senescence-associated secretory phenotype components. Transcription factor CBFβ was found to be lost in these aged cells and preferentially present in proliferating cells, indicating that CBFβ deficiency might push alveolar macrophages towards senescence (27).

Overall, current knowledge indicates differences in dysregulation of macrophage function with age that could be tissue and/or stimulus-specific. It has previously been reported (mostly using murine cells) that macrophages undergo functional decline with age, including a reduction in phagocytosis and chemotaxis (17,19). However, there is a lack of consensus as to whether, and to what extent this also occurs in primary human macrophages. Additionally, the underlying mechanisms driving these age-related changes are largely unknown. We therefore investigated the impact of ageing on primary human macrophage function, principally in monocyte-derived macrophages (MDMs), to characterise intrinsic, age-related changes to macrophage function and reveal the molecular mechanisms that underpin these changes. We found that ageing significantly reduced macrophage function and altered morphology. This age-related dysfunction and the morphological changes were replicated with knockdown of either *MYC* or *USF1* (two transcription factors with reduced expression in aged macrophages) in MDMs isolated from young donors. Finally, we identified key transcriptional targets downstream of *MYC* and *USF1* with age-related dysregulated expression that may be driving age-dependent changes in macrophage morphology and function.

## Results

### 1.1 Human monocyte-derived macrophage function is reduced with age

Monocytes were isolated from healthy sex-balanced volunteer blood aged 18-30 years (mean age 23.7 ± 1.2 years; young cohort) or > 50 years (mean age 60.5 ± 5.7 years; older cohort) (Table 1) and differentiated into monocyte-derived macrophages (MDMs). Functional assays were then performed (Fig 1) in both resting (M^0^) MDMs and cells further polarised for 24 hours with LPS and IFNγ stimulation (M^LPS+IFNγ^ MDMs). The phagocytic capacity of MDMs was assessed by the uptake of opsonised fluorescent beads (Fig 1A-B) and measured as the mean fluorescence intensity, indicative of the number of beads taken up by each cell (Fig 1C).

**Figure 1.**
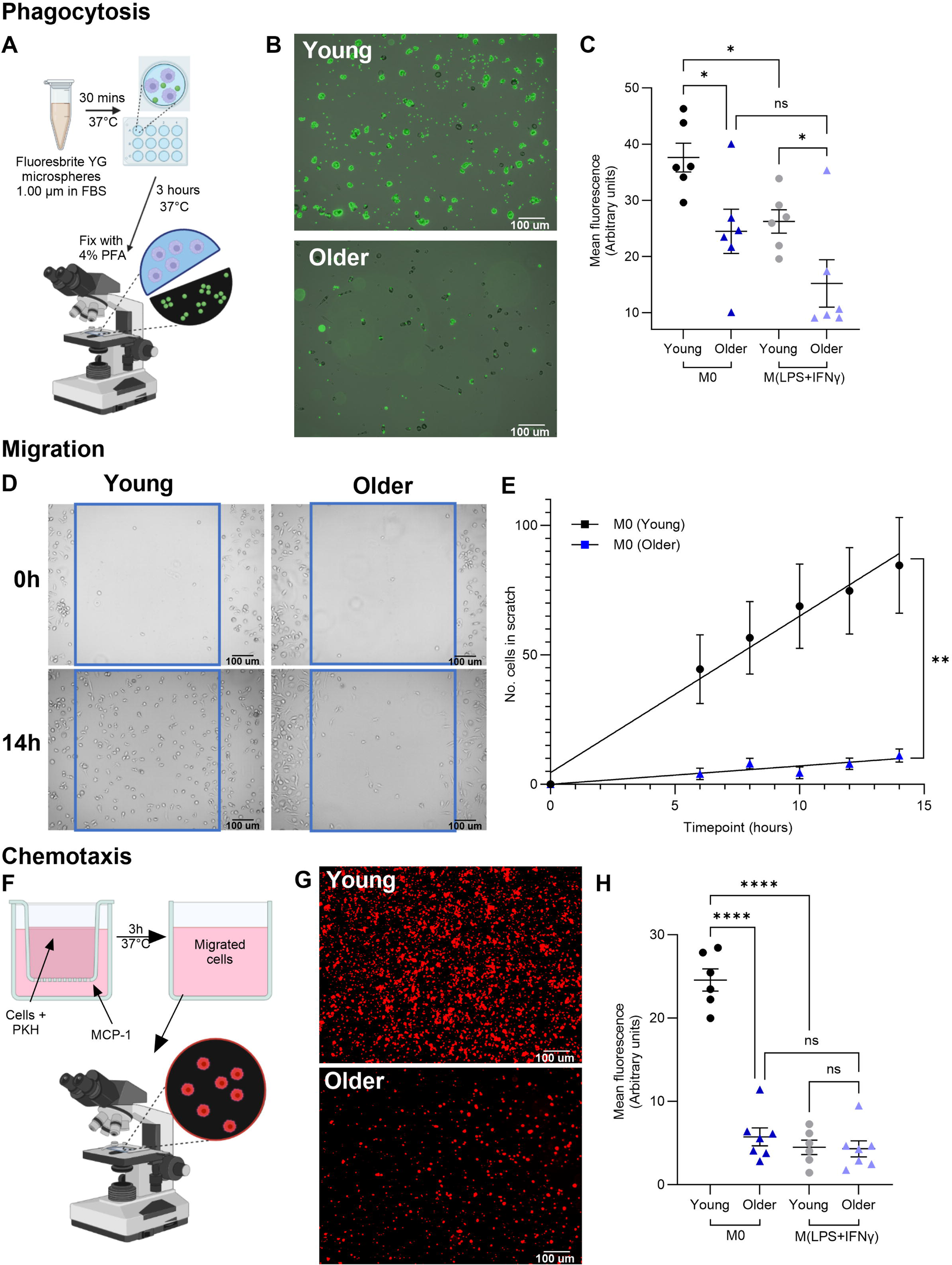
Human monocyte-derived macrophage phagocytosis and chemotaxis are reduced with age. A. Schematic diagram of phagocytosis assay design. Fluoresbrite YG microspheres 1.00 μm (Polysciences) were added to FBS and incubated at 37°C for 30 minutes before incubation for three hours with cultured MDMs. Cells were washed and fixed with 4% PFA before taking brightfield and GFP images. ImageJ “Analyze Particles” function was used to select cells in brightfield image to produce ROIs. ROIs were overlayed onto the GFP image and mean fluorescence intensity was measured. B. Representative images of GFP staining of cells from young and older subjects of GFP overlayed onto brightfield image. C. Bead uptake expressed as mean fluorescence intensity for MDMs from young and older subjects. D. Representative images of MDMs from young and older subjects returning to the scratch after 14 hours of incubation. A box was drawn around the scratch at 0 hours and replicated onto the following timepoint images from the same well to count returning cells over time. E. Number of MDMs returning to the scratch from young and older subjects. F. Schematic diagram of migration assay design. MCP-1 (5 nM) was added to the lower compartment of each well and PKH26-stained MDMs to the transwell and incubated at 37°C for 3 hours before imaging the lower cell. Mean fluorescence was measured using ImageJ for each image. G. Representative images of PKH26-stained MDMs that migrated through the transwell. H. Migrated MDMs measured as mean fluorescence intensity from young and older subjects. C,E,H. Two-way ANOVA with Tukey’s multiple comparison, * P < 0.05, **** P < 0.0001. Data are represented as mean ± SEM with each datapoint representing the mean of three fields of view taken per donor for each condition. MDMs were differentiated from human blood monocytes for 7 days in M-CSF and then either left at rest (M^0^ MDM) or further polarised for 24 hours with LPS and IFNγ (M^LPS+IFNγ^ MDMs). Young (N=6, 22-25 years), old (N=6, 54-71 years).

**Table 1.** Demographics of human donors.

| Age range | Mean age | Sex |
| --- | --- | --- |
| 18-30 years | 23.7 ± 1.2 years | F = 4, M = 2 |
| > 50 years | 60.5 ± 5.7 years | F = 3, M = 3 |

F – female; M - male

Monocyte-derived macrophages from older humans had severely reduced phagocytic capacity compared to cells isolated from young donors (Fig 1A-C). Compared to the cells from the younger subjects, phagocytosis in M^0^ MDMs from older subjects was reduced by 35% (P < 0.05) and by 42% (P < 0.05) in older M^LPS+IFNγ^ MDMs. In young cells, M^LPS+IFNγ^ MDMs showed 30% (P < 0.05) reduction in phagocytosis compared with M^0^. Older M^LPS+IFNγ^ MDMs also showed a slight reduction in phagocytosis compared with M^0^ MDMs of the same age. However, this was not significant, as mean fluorescence in older M^0^ MDMs was already reduced to levels similar to young M^LPS+IFNγ^ MDMs (mean ± SEM: 24.46±3.93 vs 26.23±2.08) (Fig 1C).

Similar to phagocytic capacity, cell migration in M^0^ MDMs, as assessed in a scratch assay (Fig 1D-E), was significantly reduced in older donors compared to young (89% less migration in macrophages from older compared to young after 14 hours migration in a scratch wound assay; P < 0.01 Two-way ANOVA). Compared to the M^0^ MDMs, migration of M^LPS+IFNγ^ MDMs in the young group was much lower (98% reduction in after 14 hours; P = 0.0021), but was comparable in both M^0^ MDMs and M^LPS+IFNγ^ MDMs in the older group (Supplementary Fig 1), illustrating the varying effects of ageing with macrophage phenotype.

To assess chemotactic migration, we used an MCP-1 transwell assay (Fig 1F). After three hours, chemotaxis of M^0^ MDMs was 77% lower in the older subjects compared with the M^0^ MDMs from the younger group. As with migration, chemotaxis of the M^LPS+IFNγ^ MDMs was significantly lower compared to M^0^ MDM population and there was no difference between the two age groups (Fig 1G-H).

Overall, these results show a reduction in M^0^ MDM function with age that was not replicated in the inflammatory polarised M^LPS+IFNγ^ cells, which had overall lower activities that resembled that seen in the M^0^ MDMs from the older subjects.

### 1.2 Six transcription factors are associated with age-related transcriptomic alterations in murine alveolar macrophages

Having established a clear phenotypic difference in aged human M^0^ MDMs (Fig 1), we set out to uncover the transcriptional networks that may drive these changes. First, we analysed a published microarray dataset from alveolar macrophages isolated from young (2-4 months) and old (22-24 months) C57BL/6J mice (14), hypothesising that transcription factors with many target genes differentially expressed with age would likely be governing the age-related changes seen in macrophage function. This was one of very few published datasets assessing transcriptional changes with age. Although alveolar macrophages are tissue resident cells with little monocyte infiltration during homeostasis, they primarily function in immune surveillance, making this dataset the most relevant to our work (24,28).

Differentially expressed genes (DEGs) with age included 213 upregulated genes (LogFC > 1.5, FDR < 0.05) and 122 downregulated genes (LogFC < −1.5, FDR < 0.05; Supplementary Fig 2, Supplementary file 1). Enrichment analysis of these genes using “ENCODE_and_ChEA_Consensus_TFs_from_ChIP-X” library in Enrichr was performed to identify the transcription factors responsible for differentially expressed transcripts (Fig 2A-B, Supplementary Fig 2). Whilst five transcription factors showed weak associations with upregulated genes, none of which reaching statistical significance (Fig 2A), six transcription factors (*Myc*, *Elf1*, *Foxm1*, *Nfyb*, *Usf1* and *Srf*) were significantly associated with the downregulated gene group (P < 0.05) (Fig 2B). Assessing expression of these transcription factors themselves between the young and aged alveolar macrophages in this dataset, (with focus on the transcription factors differentially expressed in the same direction as their target genes) showed *Myc*, *Foxm1*, *Nfyb*, *Usf1* and *Srf* were significantly downregulated and *Nfic* was upregulated with age, making them the most likely regulators to be driving age-related differential gene expression. In line with this, we found 23.8-54.2% of the putative target genes of these transcription factors that are expressed in macrophages to be dysregulated with age (Fig 2C-F).

**Figure 2.**
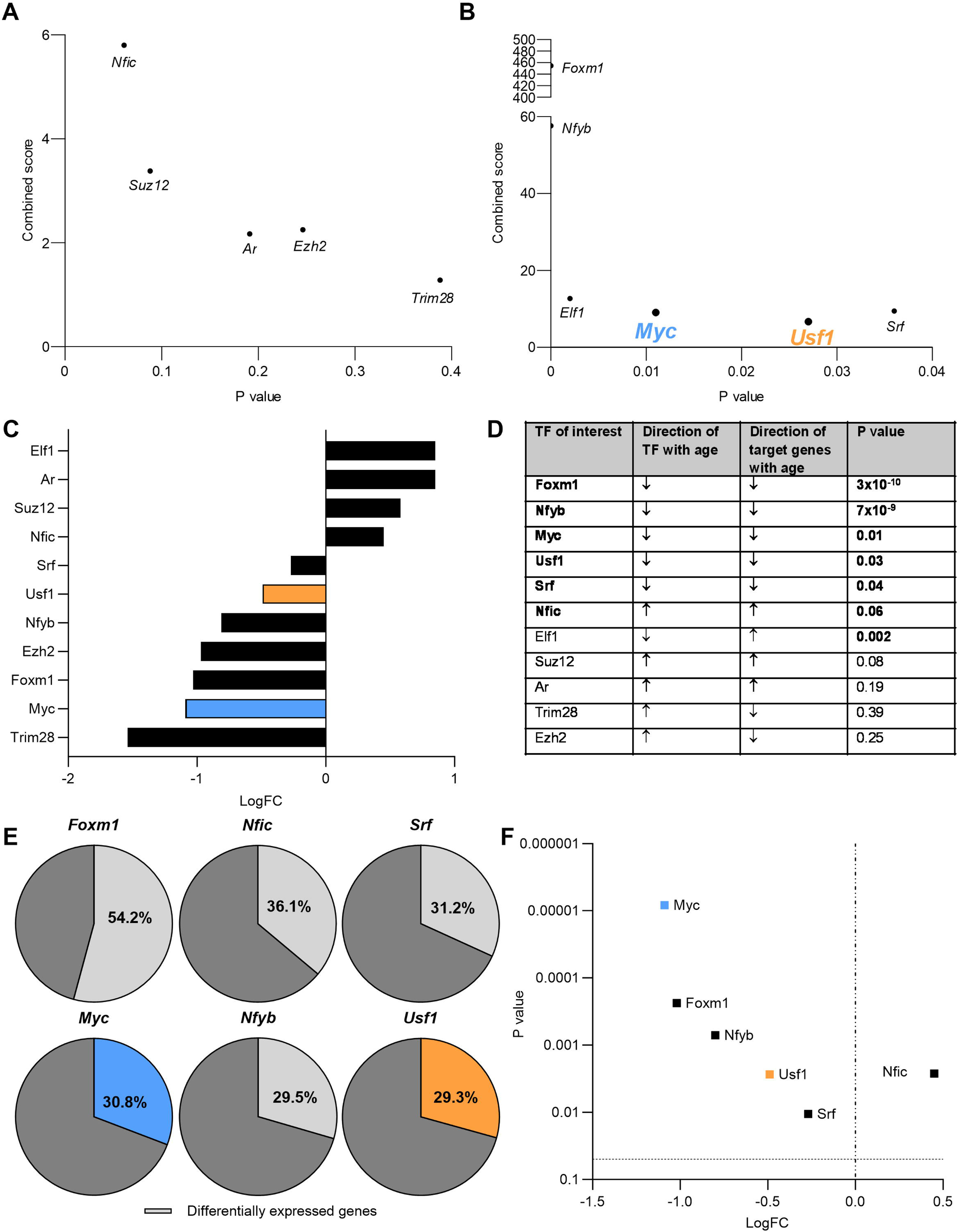
Transcription factors associated with age-related differentially expressed genes in murine alveolar macrophages. A. Transcription factors associated with upregulated genes with age. Combined score measures a combination of p value and z score to enable comparison of rankings. P value is calculated using Fisher’s exact test. B. Transcription factors associated with downregulated genes with age. Combined score measures a combination of p value and z score to enable comparison of rankings. P value is calculated using Fisher’s exact test. C. Direction and magnitude of differential expression of transcription factors associated with target gene list in old vs young. D. Ranking of transcription factors, differentially expressed with age: differentially expressed in same direction as target gene list, significant association with target gene list. E. Percentage of transcription factor target genes expressed in macrophages that are dysregulated with age. *Foxm1* – 45 differentially expressed/ of 83 targets, *Nfic* – 73/202, *Srf* – 82/ 263, *Myc* – 412/ 1336, *Nfyb* – 886/ 3004, *Usf1* – 352/1200. F. Fold change of transcription factor levels and P values adjusted for multiple-testing correction. Adjusted P value is calculated using Benjamini and Hochberg false discovery rate, (FDR < 0.05). Differentially expressed genes with age (LogFC > 1.5) were identified in GSE84901 microarray dataset comparing alveolar macrophages from young (2-4 months) and old (22-24 months) C57BL/6J mice.

### 1.3 *MYC* and *USF1* are downregulated with age in human and murine macrophages and their allelic variants are associated with monocyte percentage in human blood

To validate the six identified transcription factors as potential regulators of the observed ageing human macrophage phenotype, we compared mouse BMDMs isolated from young (2-4 months) and aged (22-24 months) C57BL/6J mice and our human cohort (Fig 3A-B).

**Figure 3.**
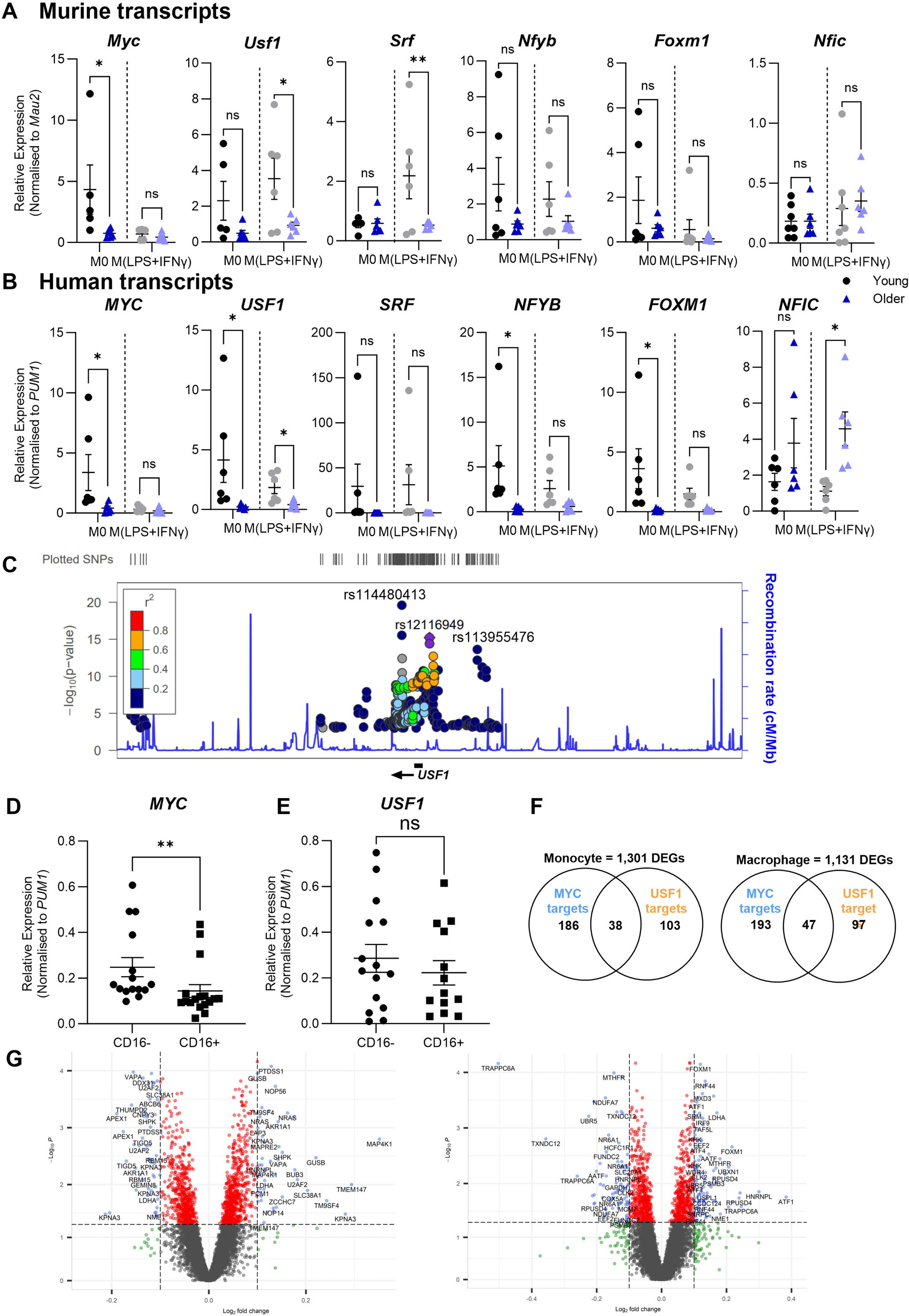
Transcription factor expression with age in murine bone marrow-derived macrophages and human monocyte-derived macrophages and genetic associations. A. Age-related changes in expression of transcription factors in bone marrow derived macrophages isolated from young (2-5 months) and aged (22-24 months) C57BL/6J mice. *Mau2* expression was used as a housekeeping control. B. Age-related changes in expression of transcription factors in human monocyte derived macrophages isolated from young (22-25 years) and older (54-71 years) healthy donors. *PUM1* expression was used as a housekeeping control. A,B. M^0^ – cells left unstimulated, M^LPS+IFNγ^ – cells stimulated with LPS and IFNγ for 24 hours. Unpaired T-test with Mann-Whitney test, N=6, * P < 0.05. C. Manhattan plot of discrete signals of eQTLs associated with *USF1* in Cardiogenic Consortium transcriptomic study monocytes. D. Relative expression of *MYC* monocyte subsets from published datasets. *PUM1* expression was used as a control to normalise all samples. Mann-Whitney test, N=15, ** P < 0.01. E. Relative expression of *USF1* monocyte subsets from published datasets. *PUM1* expression was used as a control to normalise all samples. Mann-Whitney test, N=15, ** P < 0.01. F. Association between *MYC* and *USF1* targets and differentially expressed genes with age in Cardiogenics Consortium transcriptomic study monocytes (left) and macrophages (right). G. Volcano plots of *MYC* and *USF1* targets in differentially expressed genes with age in Cardiogenic Consortium transcriptomic study monocytes (left) and macrophages (right)..

*MYC* and *USF1* showed consistent downregulation with age in both murine BMDMs (Fig 3A) and human MDMs (Fig 3B), occurring in M^0^ cells for *MYC* (while the inflammatory M^LPS+IFNγ^ phenotype had a complete lack of expression) and in both M^0^ and inflammatory stimulated M^LPS+IFNγ^ for *USF1*, although M^0^ had better overall *USF1* expression in the young human MDM population. These two transcription factors were therefore analysed further to mechanistically assess their contribution to the ageing macrophage phenotypes. Although some consistency was seen in the other assessed transcription factors compared to the murine alveolar macrophage population (*NFIC* maintained upregulation with age in human M^LPS+IFNγ^ MDMs, *NFYB* and *FOXM1* were both downregulated with age in human M^0^ MDMs), lack of consistency across both assessed cell types meant these were not taken forward.

Publicly available transcriptomic data suitable for the analysis of age-related changes in human macrophages is very limited. However, the Cardiogenics Consortium transcriptomic study dataset includes monocyte and macrophage expression profiles and genotype information from 596 donors of varying ages (42-65 years) (29–31). The macrophages in this dataset are also consistent with those in the present study that we refer to as M^0^ MDMs. To identify allelic variants of *MYC* and *USF1*, which are associated with RNA levels for these transcription factors, we analysed top and bottom age quartiles of this dataset to identify eQTLs. A total of 569 single nucleotide polymorphisms (SNPs) were associated with *USF1* and 36 SNPs was associated with *MYC* in monocytes (FDR < 0.05). A further 491 and 1 SNPs were associated with *USF1* and *MYC* in M^0^ MDMs, respectively (Supplementary file 2). All of these were cis-eQTLs and for *USF1* represented three discrete association signals (Figure 3C). Exploration of other published datasets within Phenoscanner revealed a further 9 signals (Supplementary Table 1). Sentinels for all 12 *USF1* associations were assessed for association with human traits and disease in GWAS catalogues and databases, including Phenoscanner and Open Target. The most notable genome-wide associations were found to be with blood cell traits, including blood monocyte percentage. Whilst data on age-related changes in blood monocyte counts is sparse and there is currently no consensus on whether this is altered in aged individuals, it is widely accepted that the ratio of classical (CD14^+^CD16^−^) *vs* non-classical (CD14^dim^CD16^+^) monocytes is altered, with the non-classical monocyte subset expanding during ageing. Therefore, we assessed *MYC* and *USF1* expression between classical and non-classical monocyte subsets through a combined analysis of four published datasets, equating to 15 individual samples per monocyte subset (32–35). RNA levels of *MYC* and *USF1* were normalised to the expression of *PUM1* in each datasets, a housekeeping gene found to be well conserved in our polarised and older cell populations, so that data from the individual datasets could be pooled and analysed together. We found *MYC* expression to be downregulated in CD14^dim^CD16^+^ monocytes compared with CD14^+^CD16^−^ population (Fig 3D-E), suggesting that the age-related reduction in *MYC* RNA levels could, at least in part, be due to the expansion of the low *MYC*-expressing non-classical monocyte subset. Finally, we tested whether expression of *MYC* and *USF1* target genes were altered with age in the Cardiogenic Consortium dataset by comparing top and bottom age quartiles, comprising 201 donors in each age group. Among the differentially expressed genes with age in these monocytes and M^0^ MDMs (Fig 3G, Supplementary file 2), more than 20% were putative *MYC* and *USF1* targets (Fig 3F, Supplementary file 2).

### 1.4 Loss of *MYC* or *USF1* in young human MDMs is sufficient to recapitulate an ageing macrophage phenotype

As older human macrophages displayed reduced function and both *MYC* and *USF1* were consistently downregulated with age across murine and human macrophage populations, we next investigated whether *MYC* or *USF1* expression levels are causatively linked to macrophage function. We tested this by a transient knockdown of *MYC* or *USF1* in young human MDMs and explored whether this could model the ageing macrophage phenotypes.

As shown in Fig 3A-B, both *MYC* and *USF1* were well expressed in young M^0^ populations and downregulated with age. Whereas *MYC* expression was minimal in the M^LPS+IFNγ^ phenotype, *USF1* was ubiquitously expressed in this polarised macrophage phenotype and was also downregulated with age. In contrast, expression of *MYC* and *USF1* showed no consistency in age-related dysregulation in IL-4-stimulated (M^IL-4^) macrophages, tested for completeness due to *MYC* expression being well documented in this macrophage phenotype (Supplementary Fig 3-4). Therefore, to test the sufficiency of loss of *MYC* or *USF1* to cause an old macrophage phenotype, we used siRNA gene silencing. The efficiency of *MYC* gene silencing was 97% (86.4-99.9) and of *USF1* gene silencing was 95% (81.3-99.9) in M^0^ and 83% (46.9-99.4) in M^LPS+IFNγ^, determined by quantification of *MYC* or *USF1* mRNA levels (Fig 4A). Consistent with the aged MDM phenotype, phagocytosis was reduced in both si*MYC* M^0^ and si*USF1* M^0^ compared with control M^0^. In addition, there was no further reduction in phagocytosis in si*USF1* M^LPS+IFNγ^ cells compared with M^LPS+IFNγ^ controls (Fig 4B). Similarly, migration and MCP-1 chemotaxis were reduced in both si*MYC* M^0^ and si*USF1* M^0^, compared with M^0^ control (Fig 4C-D).

**Figure 4.**
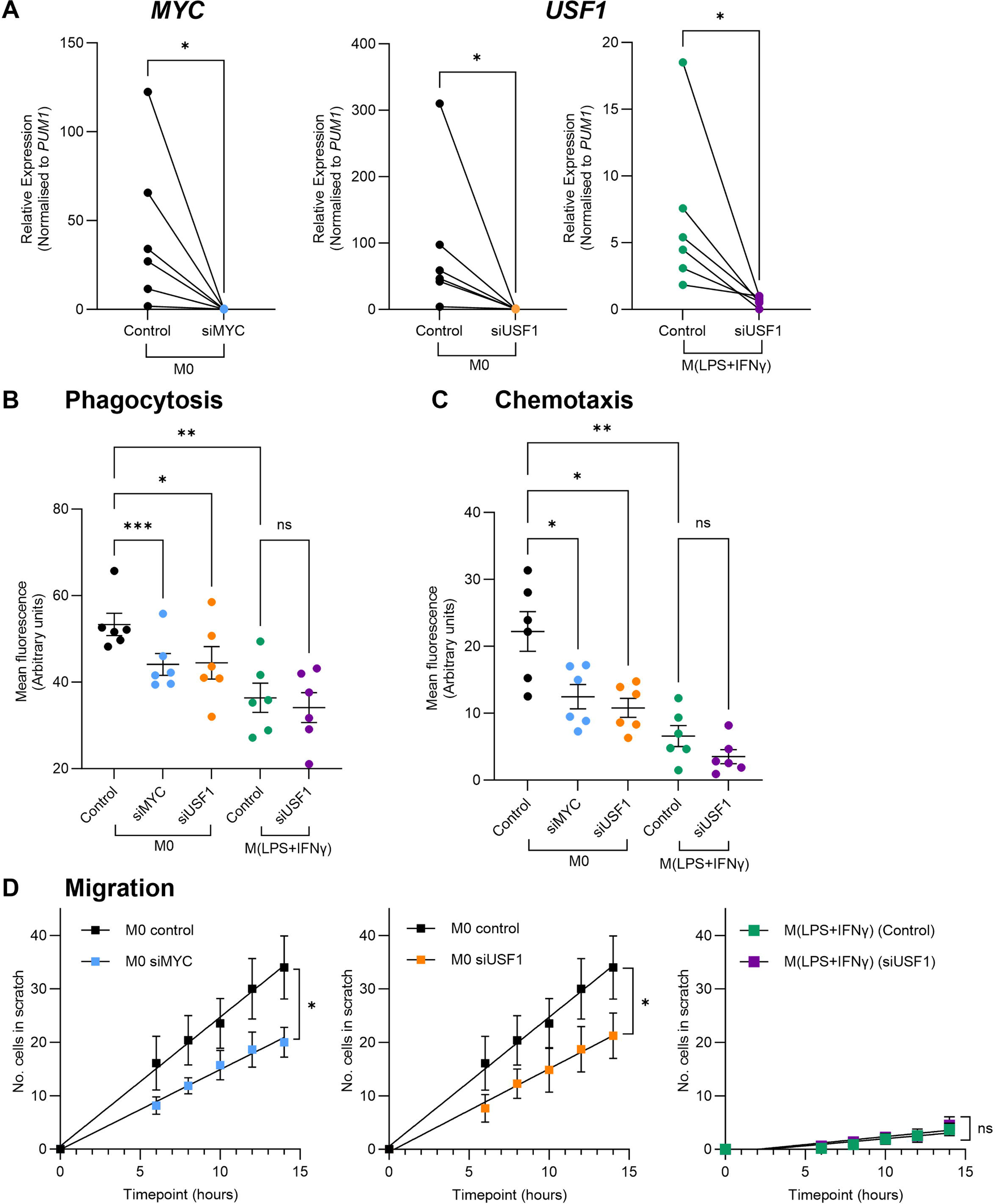
Loss of *MYC* or *USF1* in young human primary macrophages reduces phagocytosis and chemotaxis. A. Relative mRNA expression of *MYC* or *USF1* in respective siRNA knockdowns. Each datapoint represents an individual donor (N = 6). Paired T-test, * *P* < 0.05. *PUM1* expression was used as an internal control. B. Bead uptake expressed as mean fluorescence intensity from young human MDMs with loss of *MYC* or *USF1*. C. Migrated MDMs measured as mean fluorescence intensity from young human MDMs with loss of *MYC* or *USF1*. B,C. Graphs are presented as mean ± SEM. Each data point represents the mean of three images taken per donor (N=6), repeated measures one-way ANOVA with Tukey’s multiple comparison, * P < 0.05, ** P < 0.01, *** P < 0.001. D. Number of MDMs returning to the scratch from young human MDMs with loss of *MYC* or *USF1*. Graphs are presented as mean ± SEM of six donors with the mean of three images taken per donor used at each timepoint measured. Two-way ANOVA with Tukey’s multiple comparison, * P < 0.05. Control – non-targeting siRNA control, si*MYC* – *MYC*-targeting siRNA, si*USF1* – *USF1*-targeting siRNA, M^0^ – unstimulated, M^LPS+IFNγ^ – LPS and IFNγ stimulated for 24 hours after differentiation and knockdown.

### 1.5 Loss of *MYC* or *USF1* in young human MDMs reproduces the transcriptomic signature of older macrophages

To further evidence the causative role of *MYC* and *USF1* loss in the older macrophage phenotypes we had identified, we compared the transcriptomic signature of gene silenced *versus* control young macrophages (Fig 5). Principal component analysis of transcripts that were dysregulated showed a significant clustering of the knockdown conditions which extended to 5,872 DEGs (LogFC > 1, P < 0.05) between si*MYC* M^0^ and M^0^ control (Fig 5A), 6,148 DEGs between si*USF1* M^0^ and M^0^ control (Fig 5B) and 6,350 DEGs between si*USF1* M^LPS+IFNγ^ and M^LPS+IFNγ^ control (Fig 5C, Supplementary file 3).

**Figure 5.**
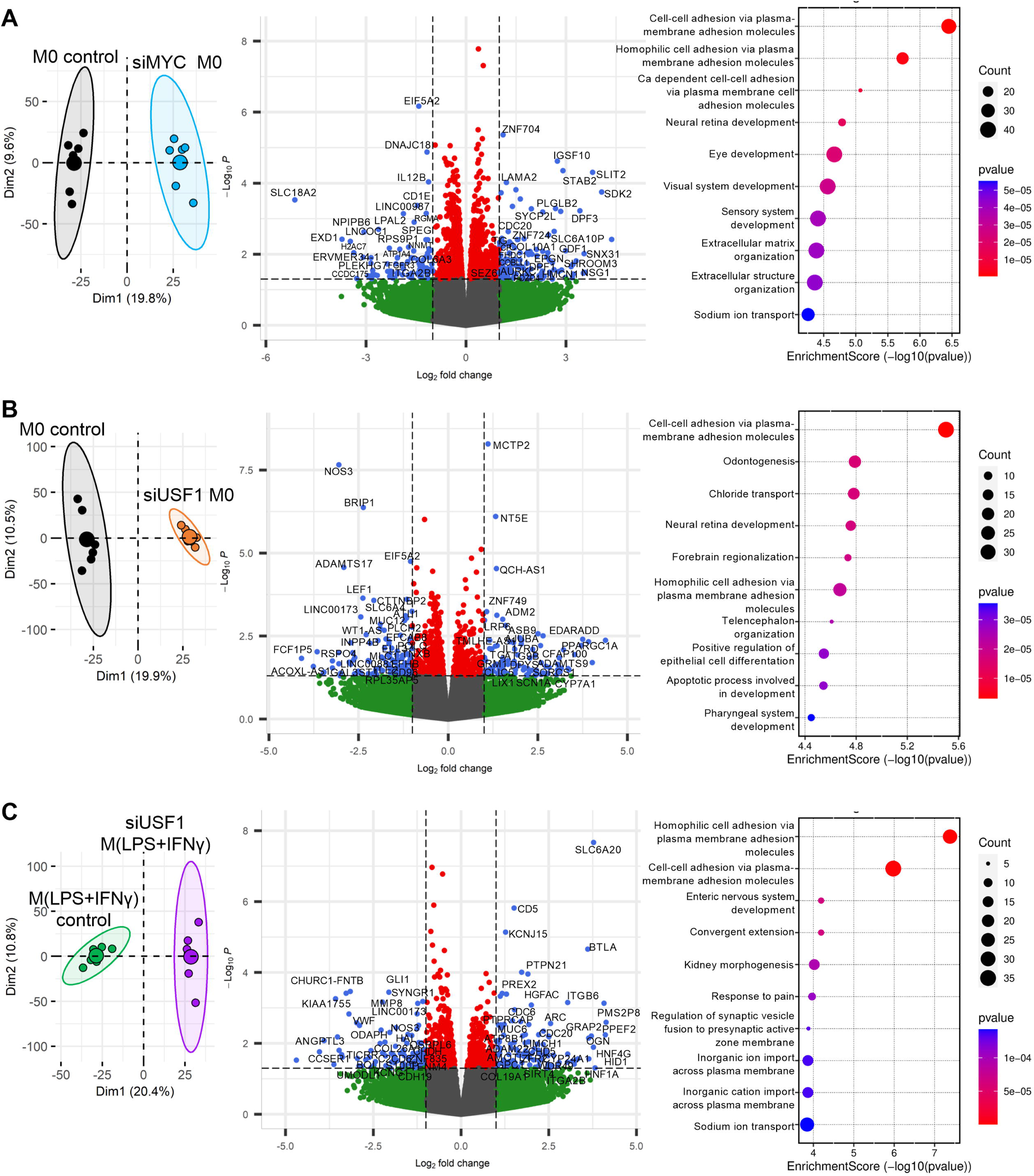
Loss of *MYC* or *USF1* produces changes in transcriptional signature associated with macrophage function. Principal component analysis of transcripts dysregulated in different conditions (left), volcano plot showing differentially expressed genes (LogFC > 1, p value < 0.05) (centre) and dot plot showing associated enriched biological processes (right) in young MDMs for si*MYC* M^0^ *vs* M^0^ control (A), si*USF1* M^0^ *vs* M^0^ control (B) and si*USF1* M^LPS+IFNγ^ *vs* M^LPS+IFNγ^ control (C).

Among the DEGs between si*MYC* M^0^ and M^0^ control, associated hallmark gene sets (36) included MYC targets, MTORC1 signalling and interferon response (Supplementary Table 2). Enriched biological processes included cell-cell adhesion, sensory development and extracellular matrix (ECM) formation (Fig 5A). Between si*USF1* M^0^ and M^0^ control, cell-cell adhesion, development, cell differentiation and apoptosis processes were enriched (Fig 5B). DEGs were also linked to hallmark gene sets including MYC targets, glycolysis and hypoxia (Supplementary Table 3). DEGs between si*USF1* M^LPS+IFNγ^ and M^LPS+IFNγ^ control were linked to hallmark gene sets such as hypoxia, angiogenesis and heme metabolism (Supplementary Table 4). Enriched biological processes included cell-cell adhesion and ion transport (Fig 5C).

To complement the analysis of human macrophages, we returned to our mouse transcriptional network dataset and performed gene set enrichment analysis (GSEA) on the DEGs from the published microarray dataset comparing young and aged alveolar mouse macrophages (14). Cell-cell adhesion and ECM components were also enriched biological processes, alongside cell cycle and DNA repair (Supplementary Table 5). Hallmarks associated with DEGs included MYC targets, G2M checkpoint and E2F targets (Supplementary Table 6).

As many DEGs were associated with similar hallmarks and biological processes between the *MYC* and *USF1* knockdowns and controls in the MDMs as well as in aged *vs* young macrophages in the mouse, we next assessed whether the same set of DEGs were also dysregulated in older human MDMs. GSEA results were used to rank DEGs by the number of times they appeared in different enriched gene sets. Leading edge analysis was then performed to sort number of appearances of each gene with its LogFC and P value (Fig 6A-C). Genes with LogFC > 1.3 and appearance in at least 5 enriched gene sets (P < 0.05), and that showed relevance to macrophage function through literature search were then analysed by RT-qPCR in human MDMs. Expression of a total of 22 genes were assessed (Fig 6D) and nine of these had reproducible differential expression in older *vs* young MDMs as was found between transcription factor knockdown *vs* control (Fig 6E-F, Supplementary Fig 5). Finally, specific analysis of *CCR2* (the MCP-1 receptor) expression, demonstrated reduced expression, providing a possible mechanism for dysregulation in MCP-1 chemotaxis (37), between si*MYC* M^0^ compared with M^0^ control and between old and young murine M^0^ BMDMs (Supplementary Fig 6).

**Figure 6.**
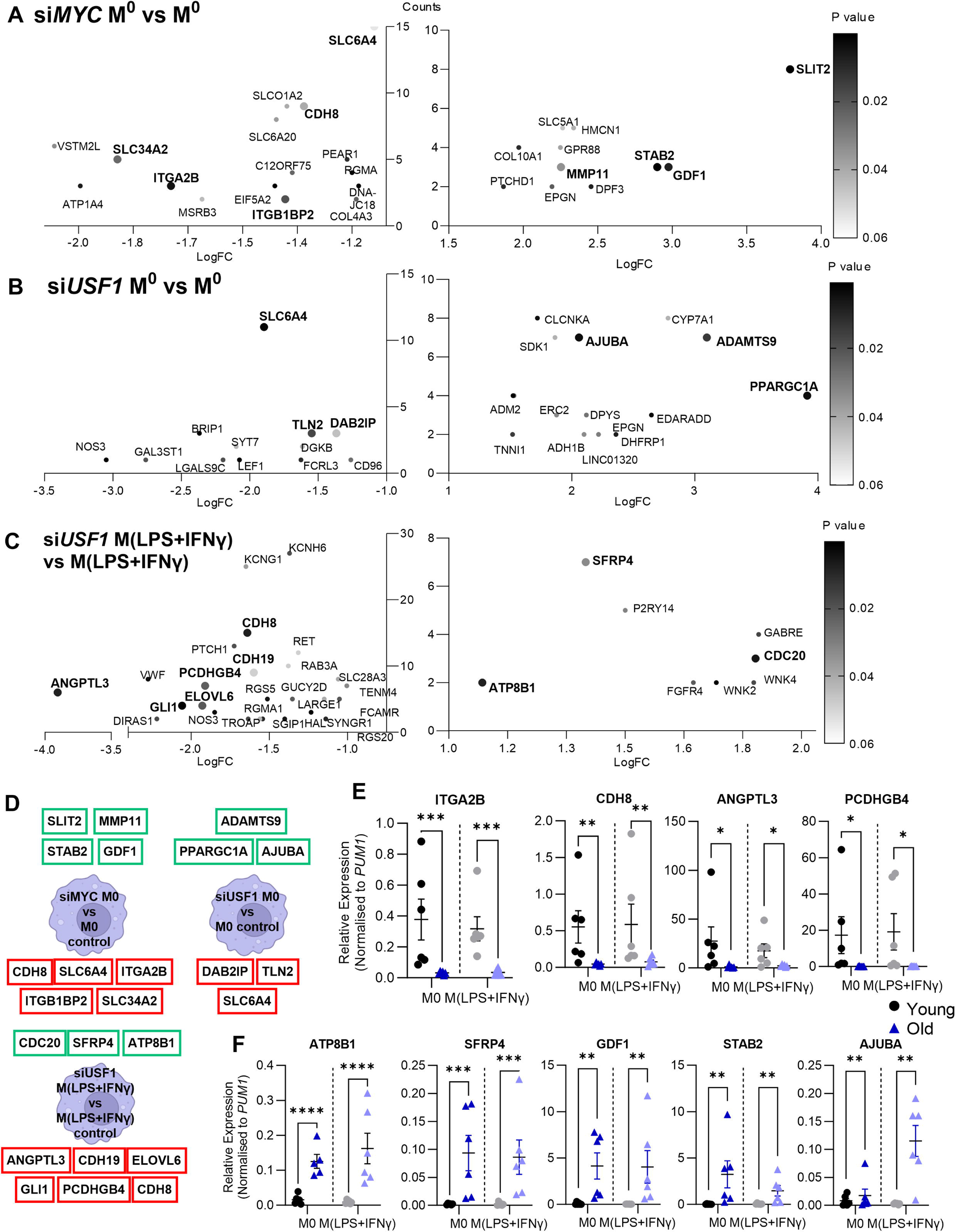
Age-related changes in gene expression are mimicked by loss of *MYC* or *USF1* in young monocyte-derived macrophages. A-C. Leading edge analysis of genes differentially expressed in si*MYC* M^0^ *vs* M^0^ control (A), si*USF1* M^0^ *vs* M^0^ control (B) or si*USF1* M^LPS+IFNγ^ *vs* M^LPS+IFNγ^ control (C), showing genes downregulated (left) and genes upregulated (right) plotted as number of enriched gene sets that the gene is associated with (y-axis) against the fold change of the gene (x-axis). Colour gradient of the dot corresponds to p value and dots of larger size were taken forward due to scoring highly in each field and association with macrophage function in literature search. D. Selected genes from each comparison taken forward for assessment in young and older human MDMs. Green: upregulated genes, red: downregulated genes E-F. Age-related changes in expression of selected genes in human monocyte-derived macrophages isolated from young (22-25 years) and older (54-71 years) healthy donors that corresponded to genes upregulated (E) or downregulated (F) in transcription factor knockdown RNA sequencing analysis. N = 6, unpaired T-test with Mann-Whitney test, * P < 0.05, ** P < 0.01, *** P < 0.001, **** P < 0.0001. *PUM1* expression was used as an internal control.

### 1.6 Age and loss of *MYC* or *USF1* in young human MDMs alter cytoskeletal structure

Having established the sufficiency of *MYC* and *USF1* loss in recapitulating the old macrophage phenotype, transcriptomic signature, and downregulation of CCR2, we used the above analysis of biological processes (Fig 4) to predict further mechanistic factors in the aging macrophage phenotype. As we found genes associated with cell-cell adhesion, ECM components and the cytoskeleton enriched in our analysis, we hypothesised that cytoskeletal and cell morphological changes would be altered in older macrophages and those with loss of *MYC* or *USF1*. From our phenotypic and transcriptional data, we predicted that older macrophages and those with loss of *MYC* or *USF1* would exhibit a more spread, adhesive appearance and reduced capability for actin motility. To test our prediction we measured cell size, circularity and F-actin (Fig 7) and found that older M^0^ MDMs were larger in size, less circular and had less F-actin content compared with young counterparts. Older M^LPS+IFNγ^ MDMs were also more elongated than young counterparts. Comparing young MDMs, M^0^s were smaller and had less F-actin content than M^LPS+IFNγ^ MDMs. Finally, older M^0^ MDMs were larger with less F-actin content than older M^LPS+IFNγ^ MDMs (Fig 7B-D). Similar findings were seen with loss of *MYC* or *USF1* in young MDMs. Compared with M^0^ control, si*MYC* M^0^ MDMs had increased size and elongation, while si*USF1* M^0^ MDMs had increased elongation as well as reduced F-actin content. Compared with M^LPS+IFNγ^ control, si*USF1* M^LPS+IFNγ^ MDMs had an increase in elongation (Fig 7E-G).

**Figure 7.**
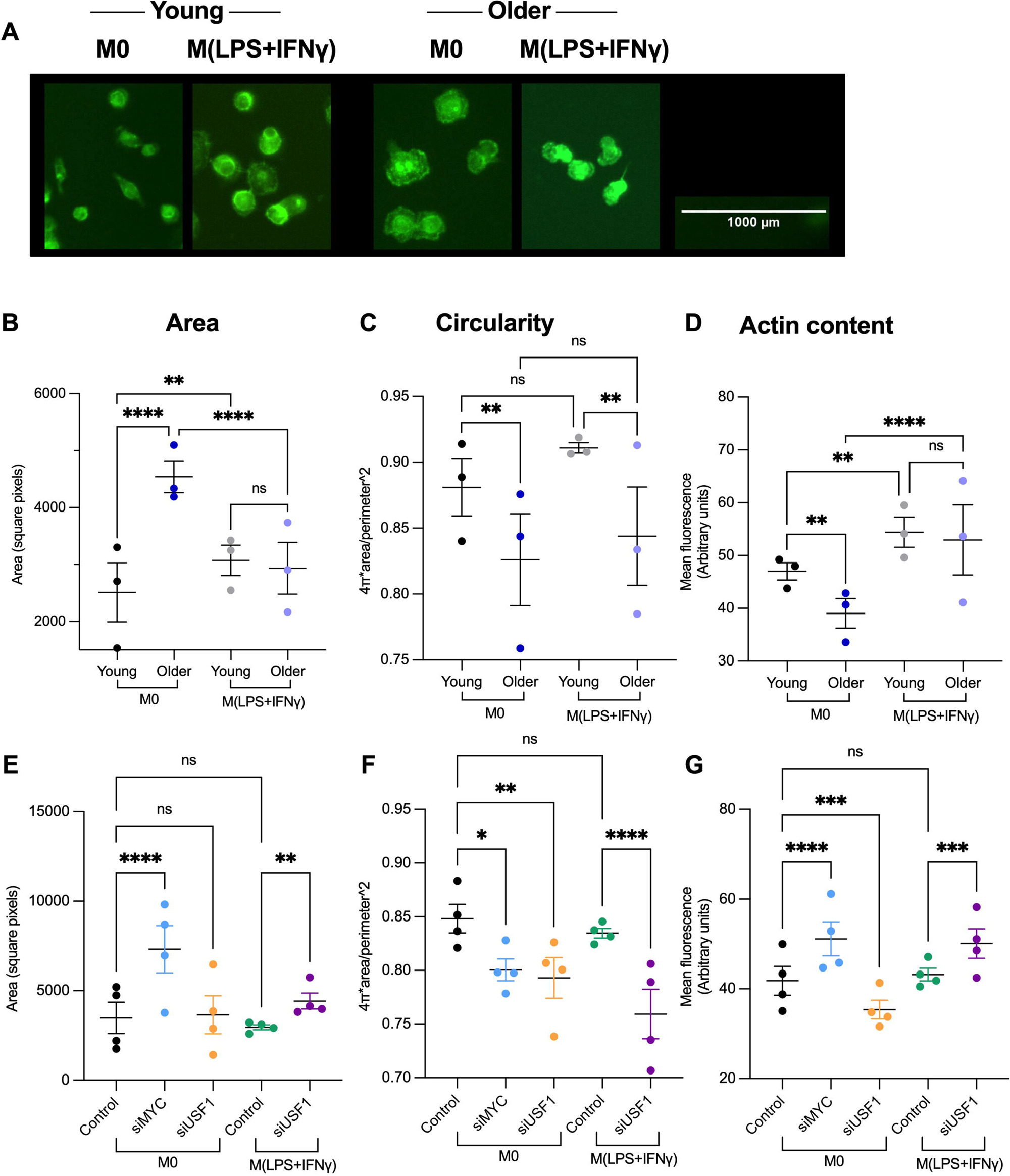
Monocyte-derived macrophage morphology and actin content change with age and with loss of *MYC* or *USF1*. A. Representative images of young and older MDMs. B-D. Morphological changes between young (N=3, 23-25 years) and older (N=3, 58-71 years) MDMs. Two-way ANOVA with Sidak’s multiple comparison, * P < 0.05, ** P < 0.01, *** P < 0.001, **** P < 0.0001 F-H. Morphological changes with loss of *MYC* or *USF1* compared with control in young MDMs (22-25 years). N=4, repeated measures one-way ANOVA with Tukey’s multiple comparison, * P < 0.05, ** P < 0.01, *** P < 0.001, **** P < 0.0001. C, F. Area by square pixel of MDMs in young and older (C) and knockdown (F) conditions. Each datapoint represents the median of four images taken per donor. D, G. Mean fluorescence intensity of MDMs stained with phalloidin to measure actin content in young and older (D) and knockdown (G) conditions. Each datapoint represents the geometric mean of four images taken per donor. E, H. Circularity (where 1 is a perfect circle and approaching 0.0 is increasingly elongated) of MDMs in young and older (E) and knockdown (H) conditions. Each data point represents the median of four images taken per donor. A minimum of 100 cells per condition were analysed across the four images. Graphs are represented as the mean ± SEM.

Thus, having identified an old macrophage phenotype in human MDMS, we had uncovered a novel *MYC*/*USF1* transcriptomic regulation of macrophage chemotaxis and phagocytosis, paired with cytoskeletal and receptor changes that can be recapitulated in young macrophages by reducing *MYC* of *USF1* RNA levels.

## Discussion

In this study, we revealed a key role for *MYC* and *USF1* transcriptional networks in age-related macrophage functional decline. We found that ageing led to a decrease in macrophage phagocytosis, migration and chemotaxis, as well as increased cell size and elongation. Loss of either *MYC* or *USF1* in young macrophages resulted in comparable functional decline and morphology, modelling the changes seen with age. We identified *MYC* and *USF1* transcripts to be downregulated with age in both murine and human macrophage populations, alongside dysregulation of transcriptional targets that may be driving age-dependent changes in function and morphology in older human macrophages.

Macrophages are key contributors to inflammaging and age-related immune decline. To date most of our understanding of macrophage ageing has been reported from mouse tissue populations, and there is a gap in functional and mechanistic understanding in human cells. Age-related changes are less well defined in monocyte-derived macrophage populations, which play a key role in age-related diseases including atherosclerosis, diabetes and fibrosis. We therefore assessed macrophage dysfunction with age in human monocyte-derived macrophages, showing both phagocytosis and chemotaxis to be reduced with age in these macrophages, that were isolated as monocytes from human blood and differentiated for 7 days in the presence of M-CSF. It is worth noting that monocyte counts between the young and older cohorts were comparable and not likely influencing these changes (10.27 ± 3.13 million vs 9.36 ± 2.9 million, respectively). While research has set out by others to assess changes in macrophage numbers with age, no real consensus has been drawn, with large tissue- and stimulus-specific variation, although it has been suggested that pro-inflammatory populations increase with age (6).

Interestingly, pro-inflammatory polarisation for 24 hours with LPS and IFNγ caused a slight but statistically significant reduction in phagocytosis compared with M^0^, both in the young and older cohorts, although this function is still thought to be necessary in the inflammatory immune response (38). M^LPS+IFNγ^ MDMs isolated from the older cohort showed the lowest phagocytic ability, with older M^0^ MDMs being similar to young M^LPS+IFNγ^ MDMs. Peritoneal macrophages have previously been assessed for phagocytosis activity after polarisation with M1-like or M2-like stimuli with comparable phagocytic activity between young and old following M2 polarisation but a reduction in phagocytosis with age in M1-like macrophages, similar to our findings (39). Fix *et al.,* also found an impairment in phagocytosis with age in pro-inflammatory BMDMs (16). In contrast to our findings in human MDMs, IFN/LPS stimulation has been found to increase phagocytosis in both young and old BMDMs compared with M^0^ (17). In our experiments, LPS+IFNγ stimulation led to a complete loss of chemotactic ability compared with M^0^ MDMs in the young. Chemotaxis in the older M^0^ MDMs was also comparable to what was seen in the young M^LPS+IFNγ^ MDMs, potentially demonstrating a chronically activated phenotype in the older M^0^ MDMs and fitting with our current understanding of inflammaging (3,5).

Both phagocytosis and chemotaxis functions have been associated with improved lifespan. Where phagocytosis has been found to be decreased with age, long-lived animals had increased phagocytosis compared with old “controls” (40). In mice, a positive correlation has been found between phagocytic ability and achieved lifespan, as well as between chemotactic capacity and achieved lifespan (41). Chemotaxis accounted for 50% of variance in lifespan, and chemotaxis and phagocytosis together were found to predict almost 70% of final achieved lifespan (42).

Systematic gene enrichment analysis of mouse alveolar transcripts identified specific transcription factors *Myc*, *Foxm1*, *Nfyb*, *Usf1* and *Srf* were downregulated and *Nfic* was upregulated with age. Consistent with this, we identified downregulated *Myc* and *Usf1* in old mouse BMDMs and in human MDMs from older donors, which led us to assess whether knockdown of *MYC* and *USF1* in young human MDMs could recapitulate an ageing phenotype. Loss of *MYC* and *USF1* significantly reduced both phagocytosis and chemotaxis in young macrophages. *MYC* is well established to play a key role in regulating the cell cycle, including proliferation, differentiation and apoptosis, as well as cell metabolism, tissue remodelling and pro- and anti-inflammatory cytokine production (43,44). It is known to be a central marker for alternative macrophage activation, controlling expression of nearly half of the genes associated with macrophage alternative activation, partly through upregulation of signal transducer and activator of transcription-6 and peroxisome proliferator-activated receptor γ signalling (45,46). *MYC* is also implicated in the growth and progression of many cancer types (47), and was shown to be the most common abnormality in malignant tumours (48). Yamanaka *et al.,* discovered that the introduction of *MYC* alongside three other transcription factors can reprogram differentiated cells back to a stem cell-like state (49); short term cyclical expression of these factors can ameliorate hallmarks of ageing and prolong lifespan (50), the first indication of *MYC*’s potential involvement in the ageing of cells. *MYC* has since been implicated in ageing, with its expression having a key role in physiological mitochondrial function (51,52), and in regulation of double-stranded DNA breaks, extensively reviewed elsewhere (53). Mitochondrial dysfunction and genomic instability are two of the hallmarks of ageing (54).

Transcription factor *USF1* has been shown to drive dysregulated cardiometabolic health (55) and cancers such as adenocarcinoma, where *USF1* expression may activate the *MYC* pathway (56), highlighting established links between these transcription factors. *USF1* responds to dietary components such as glucose, and recognises the E box CACGTG motif promoter region of target genes, many of which are involved in lipid metabolism (57). Many studied polymorphisms in *USF1* SNPs that have strong associations with plasma lipid levels, apolipoprotein E gene expression and familial hyperlipidaemia (58,59). The rs3737787 and rs2073658 genetic variants of *USF1* have significant associations with familial hyperlipidaemia, a risk factor for premature cardiovascular disease, affecting pro-inflammatory cytokine secretion and serum lipid levels (60). Further, *USF1* SNPs such as rs10908821 have an age-related association with brain legion development in Alzheimer’s Disease (61). This study has identified that *USF1* polymorphisms in macrophages and monocytes transcript from the Cardiogenic Consortium transcriptomic study and other published studies within Phenoscanner database indicate a further contribution to this ageing phenotype, since these polymorphisms are associated with decreased monocyte percentage, a known age-related change. Monocyte numbers are known to become dysregulated with age, with non-classical monocytes becoming an expanded population of the aged immune system (62,63). Aur analysis showed that *MYC* is downregulated in the non-classical compared with classical monocytes, which fits with this model. There are clear genetic associations linked to *USF1* and age-related diseases. Further, the presence of many *MYC* and *USF1* downstream targets in the differentially expressed genes of macrophages and monocytes from the approx. 600 samples of the Cardiogenic Consortium transcriptomic study cannot be understated. It has previously been shown that assessing a single gene, the transcription factor itself, is an unreliable estimation of transcription factor binding activity, especially when effect size is small, such as that of the small age range analysed in this dataset (26,64). By instead using downstream targets of the transcription factor, the challenge posed was overcome and a more reliable estimation of transcription factor activity was achieved. This analysis, therefore, further adds to our hypothesis that *MYC* and *USF1* are driving transcriptional and therefore functional changes with age.

We found that human macrophages show a profound reduction in phagocytosis, migration and chemotaxis in the older age group. We investigated the mechanistic basis for these functional changes by assessing cytoskeletal rearrangements required for coordinated phagocytosis and migration (65). Macrophage cytoskeleton is composed of cortical filamentous actin (F-actin). Resting macrophages morphology shows large actin-containing ruffes which function to survey the local environment, that are essentially precursors of phagosomes (66). We observed reduced F-actin content in older human macrophages and in young macrophages with reduced *USF1* expression, suggesting a decline in cytoskeletal function in age regulated by *USF1* which may therefore be contributing to reduced phagocytic capacity. Further, macrophage elongation has been shown to be essential for motility (67). This correlates with our finding that the young M^0^ MDMs are more motile and elongated than the M^LPS+IFNγ^ MDMs, however both phenotypes of the older MDMs had increased elongation, making this an unlikely mechanism for the reduced motility seen. Similar to our findings that older MDMs were larger than younger counterparts, older human cochlea macrophages were found to have a reduced number of processes projecting and increased cytoplasmic volume around the nucleus compared with young (68). Microglia have also been found to have larger cell bodies with increasing age, appearing as a more reactive phenotype (69). Further, aged microglia have been found to be less mobile and phagocytic with their morphology resembling dendritic cells (70,71). These aged resting cells were significantly larger than their young counterparts (71). Ageing of microglia has also been associated with an increased state of inflammation, including reduced size and longer, more extended processes, secondary branches and lamellipodia (72). Finally, Rozovsky *et al.,* found that aged microglia had an amoeboid-like structure that was not seen in young that might hinder motility (73).

As well as changes in morphology, migration requires complex adhesion interactions with other cells and the underlying extracellular matrix, a biological process that we found to be enriched by transcription factor knockdown in human MDMs and with age in murine alveolar macrophages, with integrins and cadherins being key dysregulated genes in human MDM ageing. Adherence has previously been assessed in mice with consistent findings. Peritoneal macrophages show an increase in adherence capacity with age to smooth plastic meant to resemble tissue (20), fibronectin and type 1 collagen (74). Cell adhesion pathways were also disturbed with ageing in transcriptomic analysis of microglia (75). This was comparable to the analysis performed here with six shared dysregulated genes associated with cell adhesion (PCDHGA2, AJUBA, CDH2, CXADR, TENM4, CLDN1).

Transcriptional targets of *MYC* and *USF1* that we found to be dysregulated with age suggest several further mechanisms for the ageing macrophage phenotype. *ANGPTL3* regulates lipoprotein lipase activity shown to be involved in age-related diseases such as atherosclerosis, already identified as a potential druggable target to treat this disease (76). Targeting *STAB2* ameliorates atherosclerosis by dampening inflammation (77). *ATP8B1* is associated with macrophage efferocytosis and resolution of inflammation and may be a potential therapeutic target in chronic pancreatitis (78). *SFRP4* is involved in Wnt signalling and has been implicated in skin ageing through promoting SASP, while its knockdown suppressed SASP (79). *SFRP4* is upregulated in ageing, correlating with our data, suggesting a role in macrophage chronic inflammation and ageing. Finally, *GDF1* is associated with cellular migration through Smad signalling in macrophages (80). We also considered loss of MCP-1 receptor, CCR2, as a possible mechanism for reduced migration. It has also been shown that changes in CCL2-CCR2 interaction may affect macrophage polarization towards M1, with lower CCR2 expression contributing to increased M1 polarization and higher pro-inflammatory cytokine release (37). Disruption in this interaction may therefore be pushing the older resting macrophage towards a more pro-inflammatory state. Although *CCR2* mRNA levels were downregulated with loss of *MYC* in human M^0^ MDMs compared with control and with age in murine BMDMs, human MDMs showed no statistically significant difference with age in *CCR2* expression.

In summary, we have highlighted specific functions and key regulatory factors of *MYC* and *USF1* that decline with age in human macrophages. Importantly, the older age group, showing markedly reduced phagocytosis, migration and chemotactic migration, were from healthy individuals over 50 years old with a mean age of 61 years. This average age is more than 20 years below the average life expectancy in countries with modern healthcare. The clear phenotypical changes seen in the relatively young age of our older cohort suggest that innate immune functional decline may represent an early starting point of ageing, with a culmination of factors prompting age-related disease and frailty. The underlying mechanisms of human MDM ageing are far from completely understood; the downstream functionally relevant targets of *MYC* and *USF1* that are also dysregulated with age will be key for future investigations.

### Limitations of the Study

Our studies using human monocyte-derived macrophages were from two gender-balanced healthy volunteer age groups: 18-30 years (mean age 24 ± 1.2 SD years; young cohort) or > 50 years (mean age 61 ± 5.7 SD years; older cohort). The healthy ageing are without comorbidities and frailty seen in the ageing human population, which will form more complex interesting future studies beyond the scope of this work. Human macrophage function was determined *in vitro*, since live phagocytosis and migration detection would not have been possible *in vivo*. Assessing function and gene expression under resting and specific stimulation conditions offers a simplified model compared to macrophages *in vivo*. Mouse macrophages were from male animals but these were extrapolated to human macrophage assessment from gender balanced groups.

## Acknowledgements

We thank Mark Ariaans, Jonathan Kilby and Kay Hopkinson for technical assistance. We also thank phlebotomists Salman Almalki and Saffron Foster, as well as all of the volunteers who donated blood to this research. Finally, we thank Steve Renshaw, Alison Condliffe and Colin Bingle for their editorial input. This work was supported by the Healthy Lifespan Institute, University of Sheffield.

## Author contributions

CEM, SJ, MC, VC and SH performed the experiments, data-analysis, and wrote the manuscript; AHG, SD, IS, DC, HLW and EKT designed and supervised the study and contributed to data analysis and interpretation, as well as development of the manuscript. All authors reviewed the manuscript.

## 3 STAR Methods

### RESOURCE AVAILABILITY

#### Lead contact

Further information and requests for resources and reagents should be directed to and will be fulfilled by the lead contact, Endre Kiss-Toth

#### Materials availability

This study did not generate new unique reagents.

#### Data and code availability

RNA-seq data have been deposited in GEO under accession number GSE240075 and are publicly available as of the date of publication. Microscopy data reported in this paper will be shared by the lead contact upon request. All original code has been deposited in GitHub and is publicly available as of the date of publication. DOIs are listed in the key resources table. Any additional information required to reanalyse the data reported in this paper is available from the lead contact upon request.

### EXPERIMENTAL MODEL AND STUDY PARTICIPANT DETAILS

#### Mice

C57/BL6 male mice (WT) were used at 2-5 months or 22 months of age for experiments. All animal experiments were approved by local guidelines and performed in accordance with national guidelines.

#### Human specimens

Peripheral blood mononuclear cells (PBMCs) were isolated from healthy sex-balanced donors in accordance with ethics approved by the University of Sheffield ethics committee (031330). All blood donors gave informed consent and studies were performed in accordance with the regulations of the local ethics committee.

### METHOD DETAILS

#### Human monocyte-derived macrophage isolation and culture

Blood was drawn by venepuncture and mixed with 3.8% (w/v) trisodium citrate (Na_3_C_6_O_7_). Blood was then layered onto Ficoll-Paque PLUS (GE Healthcare) and gradient centrifugation performed. Peripheral blood mononuclear cells were separated into phosphate buffer saline (PBS) containing 2mM ethylenediaminetetraacetic acid (EDTA) and red blood cells were lysed using a solution of ammonium chloride (155 mM NH_4_Cl, 10 mM KHCO_3_, 0.1 M EDTA in H_2_O). Positive selection of human monocytes was performed using magnetic CD14 microbeads (Miltenyi Biotec). Isolated monocytes were incubated in RPMI 1640 medium (Gibco) containing 10% (v/v) heat-inactivated foetal bovine serum (HI-FBS), 1% (v/v) L-Glutamine (Lonza), 1% (v/v) penicillin-streptomycin (Gibco) and 100 nM macrophage colony stimulating factor (M-CSF) (Peprotech) for 7 days at 37°C and 5% CO_2_. Differentiated monocyte-derived macrophages (MDMs) were then left unstimulated (M^0^), polarised towards an inflammatory phenotype using 100 nM lipopolysaccharide (LPS) (Enzo) and 20nM interferon-γ (IFNγ) (Peprotech) (M^LPS+IFNγ^) or polarised towards an anti-inflammatory phenotype using 20nM interleukin 4 (IL-4) (Peprotech) for a further 24 hours. Small interfering RNAs were transfected into differentiated macrophages to achieve RNA silencing prior to stimulation where applicable. Transfection reagent Fugene HD (Promega) was used following manufacturers instructions. siRNA targeting *MYC* gene (ON-TARGET plus smartpool siRNA #L-003282-02, Dharmacon) or *USF1* gene (ON-TARGET plus smartpool siRNA #L-003617, Dharmacon) was mixed at 12.5 nM with transfection reagent and Opti-MEM (Gibco). As a control, ON-TARGET plus smartpool control non-targeting (NT) siRNA (#D-001810-10-20, Dharmacon) was used at the same concentration with transfection reagent.

#### Functional assays

Functional assays were performed on human MDMs under the following conditions: young M^0^ and M^LPS+IFNγ^, older M^0^ and M^LPS+IFNγ^, and young non-targeting siRNA (M^0^ and M^LPS+IFNγ^), si*USF1*-knockdown (M^0^ and M^LPS+IFNγ^) and si*MYC*-knockdown (M^0^ only).

##### Phagocytosis assay

For phagocytosis assay, human MDMs were cultured in 12-well plates with a 4°C control and 37°C active plate. Fluoresbrite YG microspheres 1.00 μm (Polysciences) were opsonised with heat-inactivated FBS (Pan Biotech) for 30 minutes at 37°C. Five minutes before the end of incubation, control hMDMs were placed on ice. Beads were then added to each well at a concentration of 50 μL/mL and cells were incubated at 4°C or 37°C for 3 hours. After 3 hours cells were washed with PBS and fixed with 4% paraformaldehyde (PFA; Fisher Scientific) for 30 minutes at room temperature. Cells were washed again with PBS and stored in PBS at 4°C until microscopy. Images were taken using ZOE Fluorescent cell imager (BIO-RAD) at 20x zoom, with a brightfield (Gain: 5, Exposure: 340 ms, LED intensity: 59, Contrast: 20) and green channel (Gain: 40, Exposure: 500 ms, LED intensity: 43, Contrast: 0) image being taken of each field, and 3 fields being taken per temperature, condition and donor. Images were analysed using ImageJ software. The same threshold was set for each brightfield image which were then converted to mask. “Analyse particles” command was used to create regions of interest (ROIs) around each cell. ROIs were then overlayed onto GFP image and mean gray value was calculated for each cell. The geometric mean was then calculated for each field and normalised to the corresponding 4°C image. An average was then taken for each donor condition.

##### Scratch assay

To assess changes in migration, a scratch assay was performed on human MDMs cultured in 12-well plates. A scratch was made from the top to the bottom of the well, cells were washed with PBS and fresh media was added. Brightfield images (Gain: 5, Exposure: 340 ms, LED intensity: 59, Contrast: 20) were then immediately taken of the scratch to set a 0 hour threshold number of cells in the scratch. Five fields of view were taken per condition and donor. Cells were then returned to the incubator for 6 hours, from which point images were taken every 2 hours to the same specification until a 14 hour image was taken. For analysis of number of cells returning to the scratch, a box was drawn around the scratch at hour 0 and copied onto all other images taken of the same scratch. The cells inside the box were then counted and an average was taken from all fields at each timepoint for each donor. This was normalised back to hour 0 by subtracting the number of cells in the box at the start.

##### Migration assay

In order to assess ability to move towards stimuli, a transwell migration assay was performed using chemokine MCP-1. Human MDMs were cultured in 6-well plates and then collected using a cell scraper. These were centrifuged at 4000 g for 5 minutes and stained using PKH26 red fluorescent cell linker (Sigma-Aldrich) to better visualise the cell membrane, according to the manufacturer’s instructions. 1% BSA was then added to the cells to remove excess dye and these were centrifuged again at 4000 g for 5 minutes. Cells were then resuspended in 300 μL RPMI (enough to run each sample in triplicate). Using HTS Transwell 96-well permeable support, 100 μL of 1% (v/v) MCP-1 in RPMI was added to the bottom plate and 100 μL of cell suspension to the transwell insert. This was incubated for 3 hours at 37°C and 5% CO_2_. After 3 hours, the transwell insert was removed and the bottom plate was imaged using ZOE Fluorescent cell imager (BIO-RAD) at 20x zoom, with a brightfield (Gain: 5, Exposure: 340 ms, LED intensity: 59, Contrast: 20) and Red channel (Gain: 40, Exposure: 500 ms, LED intensity: 50, Contrast: 0) image being taken of each field, and 3 fields being taken per sample. ImageJ was then used to analyse each image by measuring overall mean gray value of the field and taking an average for each donor condition.

#### Microarray analysis of murine alveolar macrophages

The published dataset GSE84901 (14), was re-analysed in order to assess transcription factors driving differential expression of genes with age. The pipeline for this analysis can be found in Supplementary Fig 2 and is deposited at https://github.com/cemoss1/Moss_et_al/blob/main/GSE84901_pipeline.R. The dataset is comprised of microarray analysis of alveolar macrophages from young (2-4 months) and old (22-24 months) C57BL/6J mice. The infection-challenged samples were removed allowing for analysis of the differentially expressed genes (DEGs) as a direct result of ageing. Significant DEGs were extracted using limma package in R and defined by adjusted p-value < 0.05 and fold change >1.5 (upregulated with age) or <-1.5 (downregulated with age). A heatmap showing these DEGs can be found in Appendix Fig 1. These DEGs were then separated into two lists: genes upregulated with age and genes downregulated with age and these were used as input filed for Enrichr analysis (81–83) with ENCODE and ChEA Consensus TFs from ChIP-X library to establish transcription factors likely governing their dysregulation.

##### Further bioinformatic analysis of identified transcription factors

The transcription factors identified as governing DEGs with age were searched for in the original DEG lists and those that were differentially expressed in the same direction as their differentially expressed target genes were taken forward. The published dataset Cardiogenics Consortium transcriptomic study (29–31), comprising 596 donor MDMs (the same as the *in vitro* M^0^ MDMs in this study) and monocyte transcriptional profiles (Illumina’s Human Ref-8 Sentrix Bead Chip arrays, Illumina Inc., San Diego, CA), was re-analysed by ranking donors by age and comparing the top and bottom age quartiles (age range of 42-65) to assess age-related differentially expressed genes. It was further used to assess eQTLs present within the dataset associated with *MYC* and *USF1* in monocytes and macrophages. These eQTLs were previously identified and made available, as described in Rotival et al. (30). They were used to identify discrete signals using the locus zoom webserver and looked up in various GWAS catalogues and databases including Open Target (84,85) and Phenoscanner, both also used to find further signals in published datasets (86,87). Finally, *MYC* and *USF1* putative targets within the age-related differentially expressed genes in this dataset were assessed using the target gene library compiled on Enrichr: ENCODE and ChEA Consensus TFs from ChIP-X library. Transcription factor targets were compared against DEGs for monocytes and macrophages separately and the percentage of DEGs that were also target genes was measured.

In order to assess *MYC* and *USF1* expression in different monocyte subsets, GEO was searched for high-throughput datasets that directly compared classical and non-classical monocytes. Four datasets were found to do this (GSE2591, GSE18565, GSE16836 and GSE51997) and the GEO database was utilised to pull out raw expression values of *MYC* and *USF1* as well as *PUM1* that has been used throughout as a reference gene. Expression of *MYC* and *USF1* was then normalised to *PUM1* by dividing each donors *MYC*/*USF1* expression by the expression of *PUM1* in that monocyte subset. This method of normalisation was used as a way to compare each dataset without the need to consider bath effect.

#### Murine bone marrow-derived macrophage isolation and culture

All experiments on mouse samples were performed in accordance with UK legislation under the Animals (Scientific Procedures) Act 1996. Mouse leg bones were gifted to this research where they were not needed for other research purposes.

Bone marrow was isolated from femurs and tibias of young (2-5 months) and old (20-22 months) male C57BL/6J mice under aseptic conditions. Bones were flushed with DMEM (Gibco) using a syringe and 25-guage needle. The cell suspension was then passed through a 40 μm cell strainer and centrifuged at 500 *g* for 5 minutes. Samples obtained could then be resuspended in 90% (v/v) HI-FBS and 1-% (v/v) DMSO and frozen or immediately cultured in DMEM containing 10% (v/v) HI-FBS, 10% (v/v) L929-conditioned (M-CSF rich) media and 1% (v/v) penicillin-streptomycin (Gibco) for 5 days at 37°C and 5% CO_2_. Differentiated bone marrow-derived macrophages (BMDMs) were left unstimulated or polarised in the same way as in hMDMs.

#### RNA isolation

Cells were first lysed in BL+TG buffer and RNA isolation was performed using ReliaPrep RNA Cell Miniprep System (Promega) according to the manufacturer’s instructions. RNA was extracted from non-targeting siRNA (M^0^ and M^LPS+IFNγ^), si*USF1*-knockdown (M^0^ and M^LPS+IFNγ^) and si*MYC*-knockdown (M^0^ only) hMDMs from 6 separate donors for RNA sequencing analysis.

#### Gene expression analysis by RT-qPCR

Following RNA isolation, concentration and purity was assessed using a NanoDrop. 100 ng total RNA per reaction was transcribed to cDNA using iScript cDNA synthesis kit (BIO-RAD), according to manufacturer’s instructions. Real time qPCR was performed using CFX384 C1000 Touch Thermal Cycler (BIO-RAD) and samples were assayed using Precision PLUS SYBR-Green mastermix (Primer design). Specific genes were analysed using primers designed using NCBI BLAST tool. Primer sequences can be found in Supplementary Table 7 - 8. All assays were performed in triplicate and normalised to expression levels of housekeeping genes M^LPS+IFNγ^ (human) or Mau2 (mouse), as these were determined to be the most suitable across young and old samples. Relative transcript levels were analysed using 2^−ΔCt^ method.

#### RNA sequencing analysis

Total RNA was isolated as previously described. An indexed pair–end sequencing run of 2 × 51 cycles on Illumina HiSeq 2000 was performed on all samples by Novogene. FASTQ files were provided which were mapped onto build 38 of the human genome using STAR (88). Mapped transcripts were then converted to genes using tximportData in R. Using edgeR, counts per million were computed and used for principle component analysis (PCA). Pairwise differential gene expression analyses for each of the comparisons were performed using DESeq2 in R with non-targeting siRNA M^0^ or M^LPS+IFNγ^ control used as the reference for gene knockdown in M^0^ or M^LPS+IFNγ^, respectively. DEGs were identified using False Discovery Rate (FDR) of < 5% and fold change > 1, when compared to reference. Gene set enrichment analysis (GSEA) (89,90), Enrichr (81–83) and GOrilla (91,92) analysis were performed on the DEG lists to output associated pathways.

#### Cytoskeleton staining and analysis

Following monocyte isolation from human blood, cells were cultured onto 13 mm coverslips in 24-well plates and left to differentiate as above. hMDMs were fixed with 4% (v/v) formaldehyde for 10 minutes at room temperature and then maintained in PBS at 4°C. At this point all samples were blinded so that the microscopy and analysis were performed without knowledge of condition or donor being assessed. Coverslips were then removed onto parafilm and cells were washed with PBS then quenched with ammonium chloride for 10 minutes before being washed again. Cells were permeabilised with 0.1% (v/v) Triton-X diluted in PBS for 4 minutes. Coverslips were then stained with Fluorescein Isothiocyanate labelled Phalloidin (Sigma-Aldrich) for 20 minutes and washed with PBS and then water. Coverslips were mounted onto slides using Mowiol and kept in the dark for at least 24 hours to dry and then until imaging. Images were taken using a microscope at 20x and 60x magnification. At 60x magnification oil immersion was used. Four fields of view per condition per donor were taken and analysed using ImageJ software. Each cell was manually drawn around and then area, circularity (shape descriptors) and mean gray value were measured. A circularity score of 1.0 indicates a perfect circle whereas values approaching 0.0 indicate an increasingly elongated shape. The median for area and circularity and geometric mean for mean gray value were then calculated for each image and an average was taken. PCA was used to analyse clustering of the cells according to size, circularity and fluorescence (Supplementary Fig 7).

### QUANTIFICATION AND STATISTICAL ANALYSIS

GraphPad Prism 9 software was used to generate all statistical analysis and graphs. N numbers and statistical analysis are stated for each experiment in the figure legends including the statistical test used, exact value of n and what n represents, definition of centre and dispersion and precision measures. Differences were considered to be significant when P values were < 0.05. Donors used in RNA sequencing were blinded prior to analysis. All macrophage samples used for microscopy were also blinded prior to imaging and analysis to prevent bias. Unless otherwise stated, six biological replicates were used for all experiments with each having three technical replicates. N = 6 donors for each condition was used due to this being previously shown to give statistical power. Where possible, when comparing the same donor under different conditions, paired analysis was performed. ANOVA was used for time course experiments (Fig 1E, Fig 4D).

### KEY RESOURCES TABLE

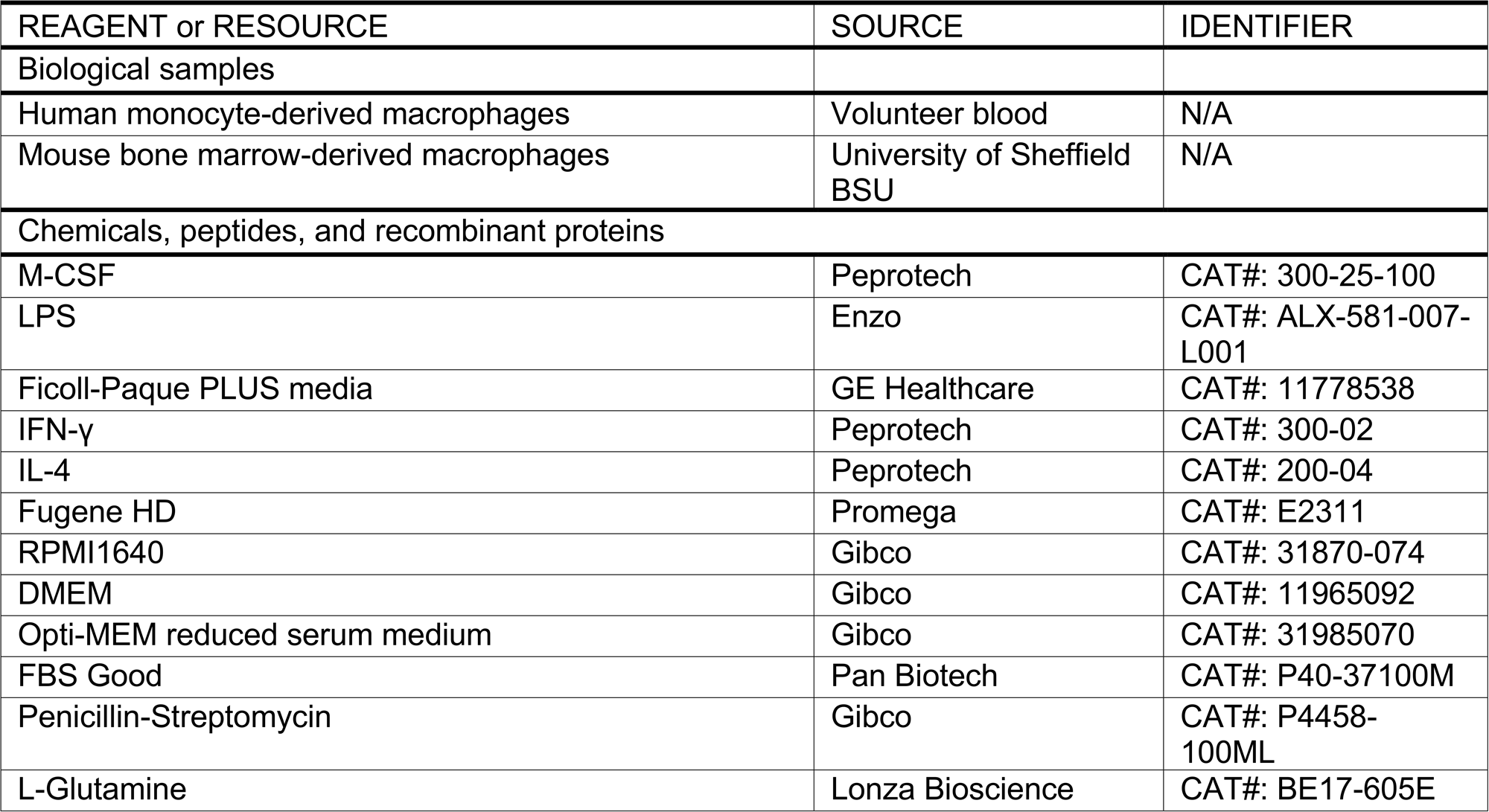
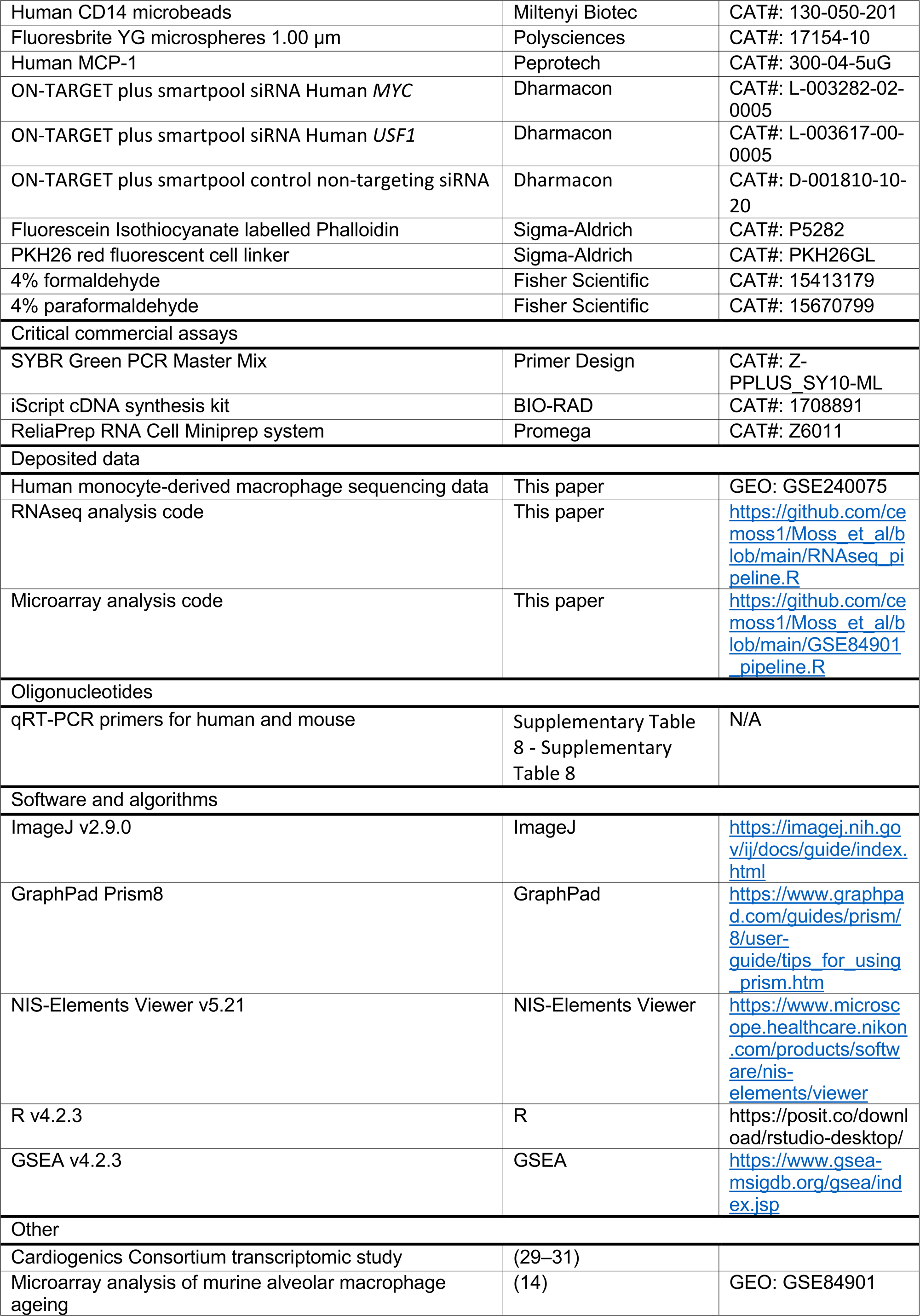

| REAGENT or RESOURCE | SOURCE | IDENTIFIER |
| --- | --- | --- |
| Biological samples |  |  |
| Human monocyte-derived macrophages | Volunteer blood | N/A |
| Mouse bone marrow-derived macrophages | University of Sheffield BSU | N/A |
| Chemicals, peptides, and recombinant proteins |  |  |
| M-CSF | Peprtech | CAT#: 300-25-100 |
| LPS | Enzo | CAT#: ALX-581-007-L001 |
| Ficoll-Paque PLUS media | GE Healthcare | CAT#: 11778538 |
| IFN-γ | Peprtech | CAT#: 300-02 |
| IL-4 | Peprtech | CAT#: 200-04 |
| Fugene HD | Promega | CAT#: E2311 |
| RPMI1640 | Gibco | CAT#: 31870-074 |
| DMEM | Gibco | CAT#: 11965092 |
| Opti-MEM reduced serum medium | Gibco | CAT#: 31985070 |
| FBS Good | Pan Biotech | CAT#: P40-37100M |
| Penicillin-Streptomycin | Gibco | CAT#: P4458-100ML |
| L-Glutamine | Lonza Bioscience | CAT#: BE17-605E |
| Human CD14 microbeads | Miltenyi Biotec | CAT#: 130-050-201 |
| Fluoresbrite YG microspheres 1.00 μm | Polysciences | CAT#: 17154-10 |
| Human MCP-1 | Peptotech | CAT#: 300-04-5uG |
| ON-TARGET plus smartpool siRNA Human *MYC* | Dharmacon | CAT#: L-003282-02-0005 |
| ON-TARGET plus smartpool siRNA Human *USF1* | Dharmacon | CAT#: L-003617-00-0005 |
| ON-TARGET plus smartpool control non-targeting siRNA | Dharmacon | CAT#: D-001810-10-20 |
| Fluorescein Isothiocyanate labelled Phalloidin | Sigma-Aldrich | CAT#: P5282 |
| PKH26 red fluorescent cell linker | Sigma-Aldrich | CAT#: PKH26GL |
| 4% formaldehyde | Fisher Scientific | CAT#: 15413179 |
| 4% paraformaldehyde | Fisher Scientific | CAT#: 15670799 |
| Critical commercial assays |  |  |
| SYBR Green PCR Master Mix | Primer Design | CAT#: Z-PPLUS_SY10-ML |
| iScript cDNA synthesis kit | BIO-RAD | CAT#: 1708891 |
| ReliaPrep RNA Cell Miniprep system | Promega | CAT#: Z6011 |
| Deposited data |  |  |
| Human monocyte-derived macrophage sequencing data | This paper | GEO: GSE240075 |
| RNAseq analysis code | This paper | https://github.com/ce-moss1/Moss_et_al/blob/main/RNAseq_pipeline.R |
| Microarray analysis code | This paper | https://github.com/ce-moss1/Moss_et_al/blob/main/GSE84901_pipeline.R |
| Oligonucleotides |  |  |
| qRT-PCR primers for human and mouse | Supplementary Table 8 - Supplementary Table 8 | N/A |
| Software and algorithms |  |  |
| ImageJ v2.9.0 | ImageJ | https://imagej.nih.gov/ij/docs/guide/index.html |
| GraphPad Prism8 | GraphPad | https://www.graphpad.com/guides/prism/8/user-guide/tips_for_using_prism.htm |
| NIS-Elements Viewer v5.21 | NIS-Elements Viewer | https://www.microscope.healthcare.nikon.com/products/software/nis-elements/viewer |
| R v4.2.3 | R | https://posit.co/download/rstudio-desktop/ |
| GSEA v4.2.3 | GSEA | https://www.gsea-msigdb.org/gsea/index.jsp |
| Other |  |  |
| Cardiogenics Consortium transcriptomic study | (29–31) |  |
| Microarray analysis of murine alveolar macrophage ageing | (14) | GEO: GSE84901 |

