## Supplementary figures and tables for "Ageing-related defects in macrophage function are driven by *MYC* and *USF1* transcriptional programmes"

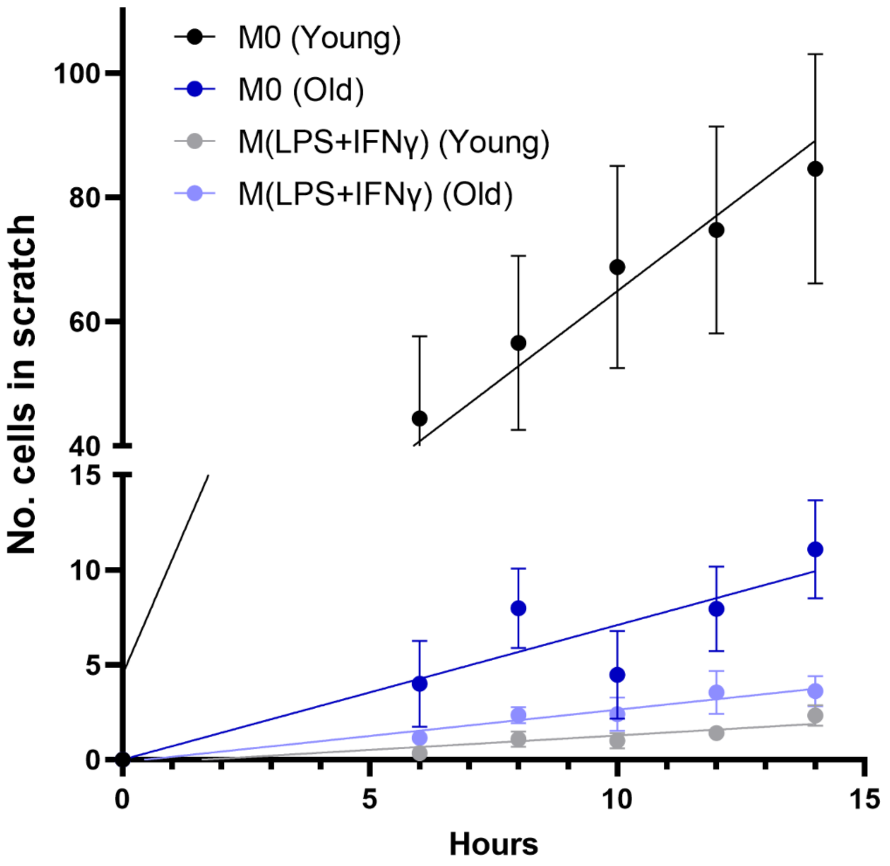


**Supplementary Fig 1 – Motility of M$^{LPS+IFN\gamma}$ polarised human monocyte-derived macrophages**

Number of cells returning to the scratch for young and old human MDMs. A line of best fit was added and area under curve was measured. All cells were imaged using ZOE fluorescent imager (BIO-RAD) with three fields of view taken per donor for each condition. Data are represented as mean ± SEM with each datapoint representing the mean of six donors, three images taken per donor. Young (N=6, 22-25 years), old (N=6, 54-71 years), M$^0$ (unstimulated), M$^{LPS+IFN\gamma}$ (LPS+IFNγ stimulated).

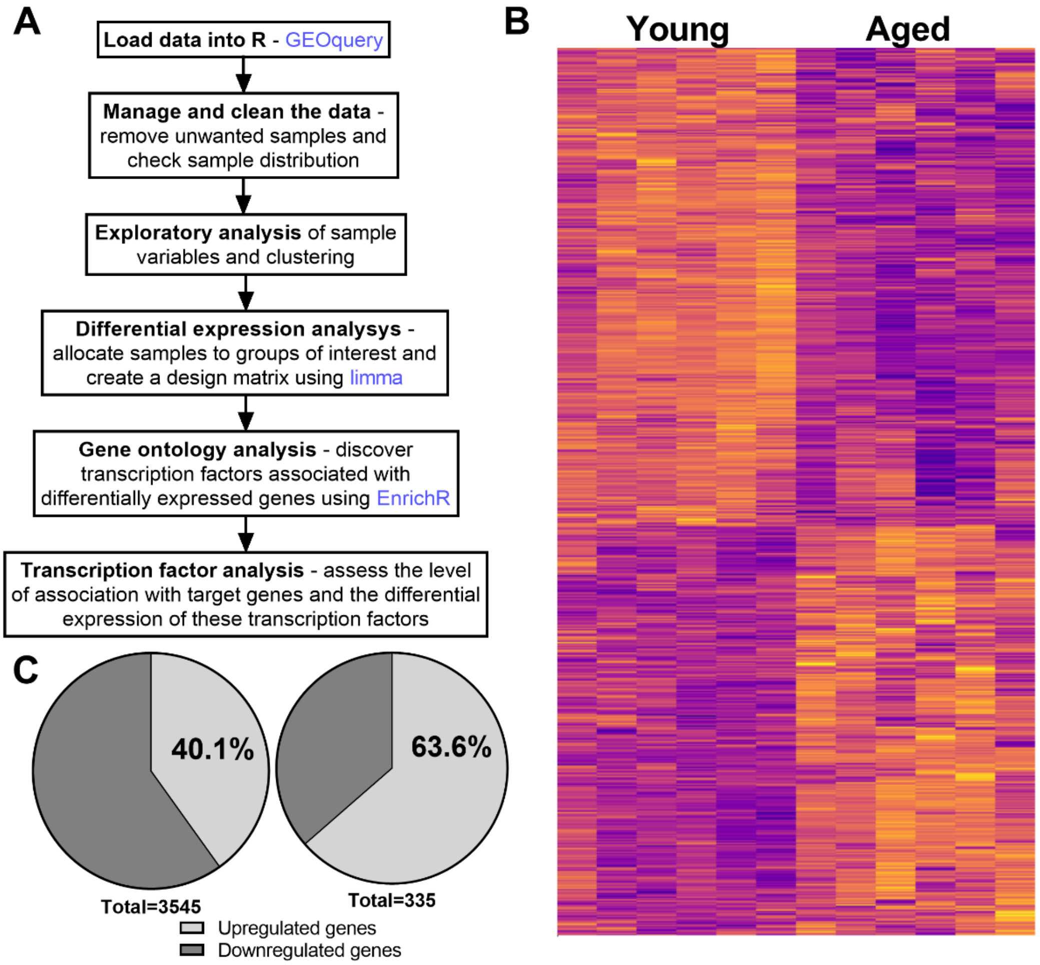


**Supplementary Fig 2 – Age-related differentially expressed genes in mouse alveolar macrophages**

A. Main steps in the pipeline for published microarray dataset GSE84901 analysis. Boxes represent the overall objectives, blue text shows the packages used.

B. A total of 3,545 genes were differentially expressed with age LogFC > 1, p-value < 0.05.

C. Of these differentially expressed genes, 1,422 (40.1%) were upregulated and 2,123 (59.9%) were downregulated with age. Increasing the threshold of differential expression to LogFC > 1.5 and p-value < 0.05, 213 genes (63.3%) were upregulated and 122 genes (36.4%) were downregulated with age.

**Supplementary table 1 – *USF1* and *MYC* discrete signals from Phenoscanner**

Data for 9 discrete ($r^2$<0.3) eQTL signals for *USF1* and 1 for *MYC* were extracted from Phenoscanner. All eQTL data is from whole blood. SNP marker (rsid), chromosome and position (hg38), effect allele (EA) and frequency (EAF, Eur) is shown alongside position in relation to the gene, source study, P-value and the direction of effect associated with the effect allele.

| Gene | Signal | SNP | Chromosome: position | EA | EAF | Position | Study | P | Direction |
|---|---|---|---|---|---|---|---|---|---|
| *USF1* | 1 | rs147573079 | chr1:161039365 | A | 0.0129 | 3_prime_UTR | eQTLGen | 3.06E-11 | - |
| | | rs75089506 | chr1:161044519 | A | 0.0129 | intron | eQTLGen | 2.99E-11 | - |
| *USF1* | 2 | rs3737787 | chr1:161039733 | A | 0.2942 | 3_prime_UTR | BIOSQTL | 6.65E-46 | + |
| | | rs2073658 | chr1:161040972 | T | 0.2942 | intron | BIOSQTL | 5.35E-46 | + |
| | | rs2073656 | chr1:161041565 | C | 0.2942 | intron | BIOSQTL | 5.35E-46 | + |
| | | rs2073655 | chr1:161042800 | A | 0.2942 | intron | BIOSQTL | 5.35E-46 | + |
| *USF1* | 3 | rs2516841 | chr1:161040984 | A | 0.2386 | intron | eQTLGen | 2.1E-285 | - |
| | | rs2774276 | chr1:161041926 | C | 0.7614 | intron | eQTLGen | 2.5E-286 | + |
| | | rs2073657 | chr1:161041001 | T | 0.6282 | intron | BIOSQTL | 1.1E-203 | + |
| | | rs2516839 | chr1:161043331 | T | 0.6292 | 5_prime_UTR | BIOSQTL | 7.6E-207 | + |
| | | rs2774273 | chr1:161044195 | C | 0.6292 | intron | BIOSQTL | 7.6E-207 | + |
| | | rs2516837 | chr1:161044937 | A | 0.3708 | 5_prime_UTR | BIOSQTL | 2.4E-207 | - |
| *USF1* | 4 | rs17221763 | chr1:161041419 | A | 0.0219 | splice_region | eQTLGen | 5.17E-35 | - |
| | | rs17175575 | chr1:161041525 | T | 0.0229 | intron | eQTLGen | 4.09E-36 | - |
| *USF1* | 5 | rs2516840 | chr1:161041527 | A | 0.2694 | intron | eQTLGen | 4.9E-119 | - |
| *USF1* | 6 | rs2073653 | chr1:161042970 | T | 0.8678 | intron | eQTLGen | 1.1E-122 | + |
| | | rs6686076 | chr1:161043522 | T | 0.8678 | intron | eQTLGen | 8.4E-123 | + |
| | | rs6427572 | chr1:161043813 | A | 0.1332 | intron | eQTLGen | 1.1E-112 | - |
| *USF1* | 7 | rs1556260 | chr1:161044656 | T | 0.1322 | intron | eQTLGen | 1.2E-122 | - |
| | | rs1556259 | chr1:161044859 | A | 0.8678 | intron | eQTLGen | 9.6E-123 | + |
| *USF1* | 8 | rs2516838 | chr1:161044580 | C | 0.666 | intron | BIOSQTL | 5.46E-55 | - |
| *USF1* | 9 | rs149397699 | chr1:161044634 | T | 0.9791 | intron | eQTLGen | 7.4E-26 | + |
| *MYC* | 1 | rs2070583 | chr8:127741008 | A | 0.9821 | 3_prime_UTR | Joehanes R | 2.09E-08 | - |

Source study: **eQTLGen**: Võsa, U., Claringbould, A., Westra, HJ. et al. Large-scale cis- and trans-eQTL analyses identify thousands of genetic loci and polygenic scores that regulate blood gene expression. Nat Genet 53, 1300–1310 (2021). https://doi.org/10.1038/s41588-021-00913-z
**BIOSQTL**: Zhernakova, D. V., Deelen, P., Vermaat, M., van Iterson, M., van Galen, M., Arindrarto, W., van 't Hof, P., Mei, H., van Dijk, F., Westra, H. J., et al (2017). Identification of context-dependent expression quantitative trait loci in whole blood. Nature genetics, 49(1), 139–145. https://doi.org/10.1038/ng.3737
**Joehanes R**: Joehanes, R., Zhang, X., Huan, T. et al. Integrated genome-wide analysis of expression quantitative trait loci aids interpretation of genomic association studies. Genome Biol 18, 16 (2017). https://doi.org/10.1186/s13059-016-1142-6

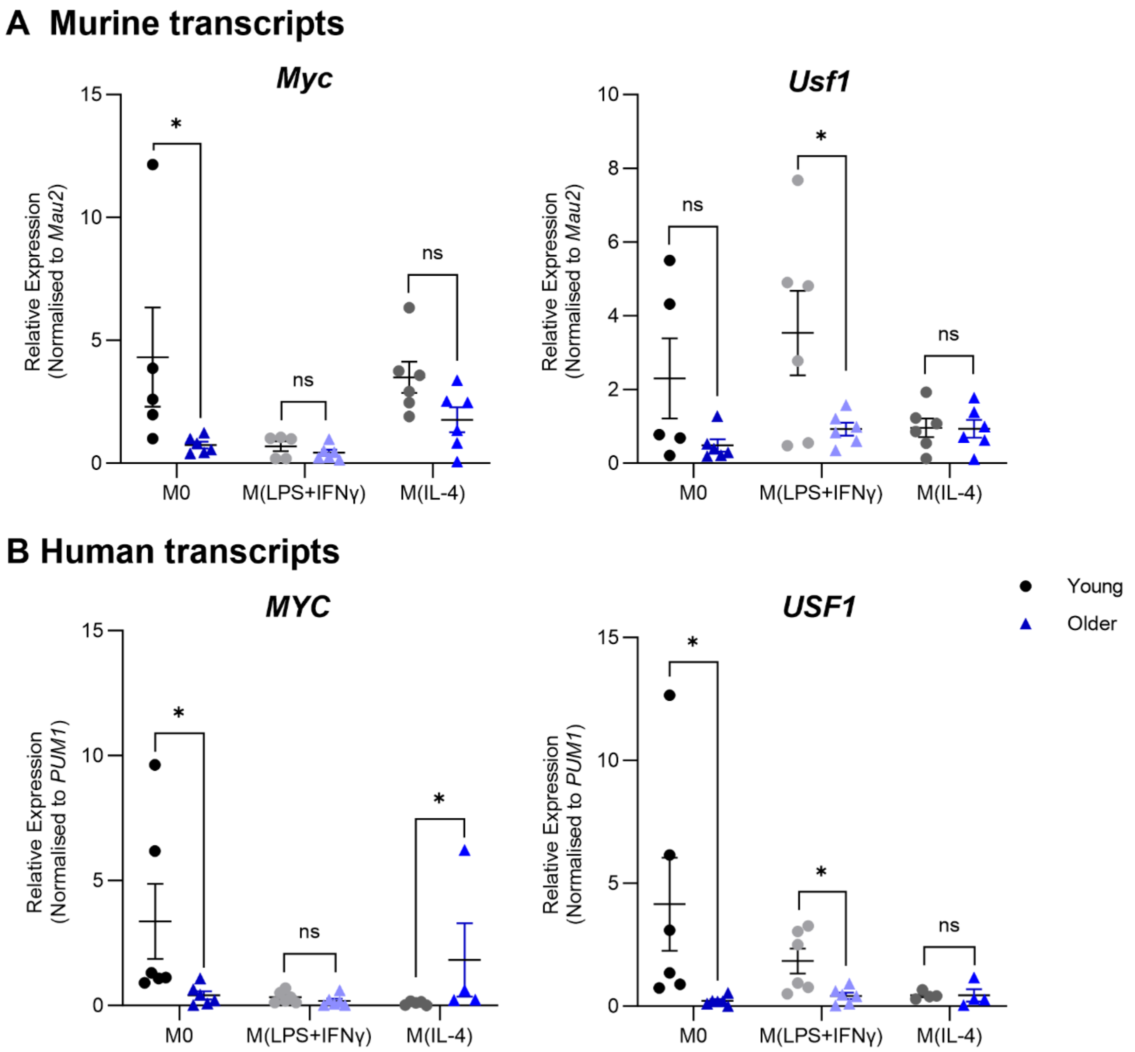


**Supplementary Fig 3 - Expression of *MYC* and *USF1* with age in murine bone marrow-derived macrophages and human monocyte-derived macrophages polarised towards different phenotypes**

A. Age-related changes in *Myc* and *Usf1* expression in bone marrow derived macrophages isolated from young (2-5 months) and aged (22-24 months) C57BL/6J mice. *Mau2* expression was used as an internal control.

B. Age-related changes in *MYC* and *USF1* expression in human monocyte derived macrophages isolated from young (22-25 years) and older (54-71 years) healthy donors. *PUM1* expression was used as an internal control.

A,B. $M^0$ – cells left unstimulated, $M^{LPS+IFN\gamma}$ – cells stimulated with LPS and IFNγ for 24 hours, $M^{IL\text{-}4}$ – cells stimulated with IL-4 for 24 hours. Unpaired T-test with Mann-Whitney test, N=6, * $P < 0.05$.

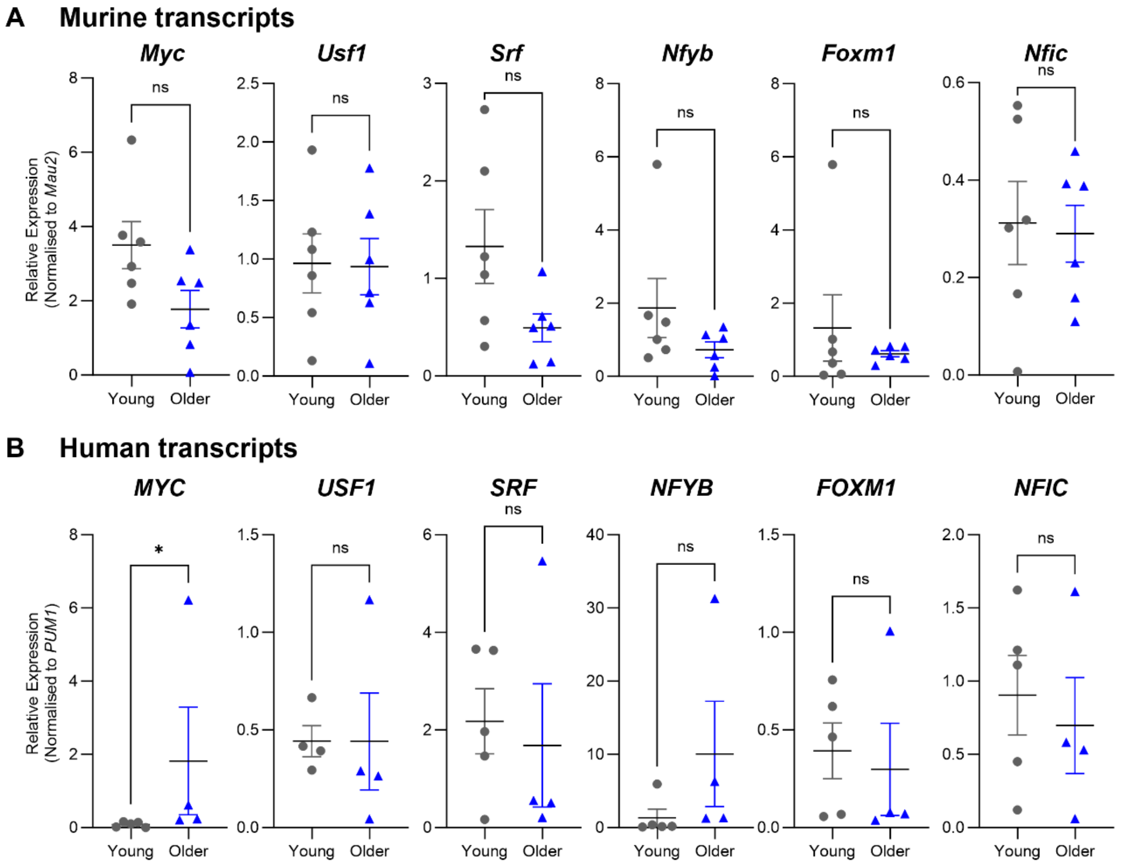


**Supplementary Fig 4 – Transcription factor expression with age in IL-4-stimulated mouse BMDMs and human MDMs**

A. Transcription factor expression by RT-qPCR in BMDMs isolated from young (2-5 months) and old (22-24 months) C57BL/6J mice. BMDMs were differentiated for 5 days with M-CSF and stimulated with IL-4 for a further 24 hours. Unpaired T-test with Mann-Whitney test, N=6.

B. Transcription factor expression by RT-qPCR in hMDMs isolated from young (22-25 years) and old (54-71 years) healthy donors. Human MDMs were differentiated for 7 days with M-CSF and stimulated with IL-4 for a further 24 hours. Unpaired T-test with Mann-Whitney test, N=4-5, * P < 0.05.

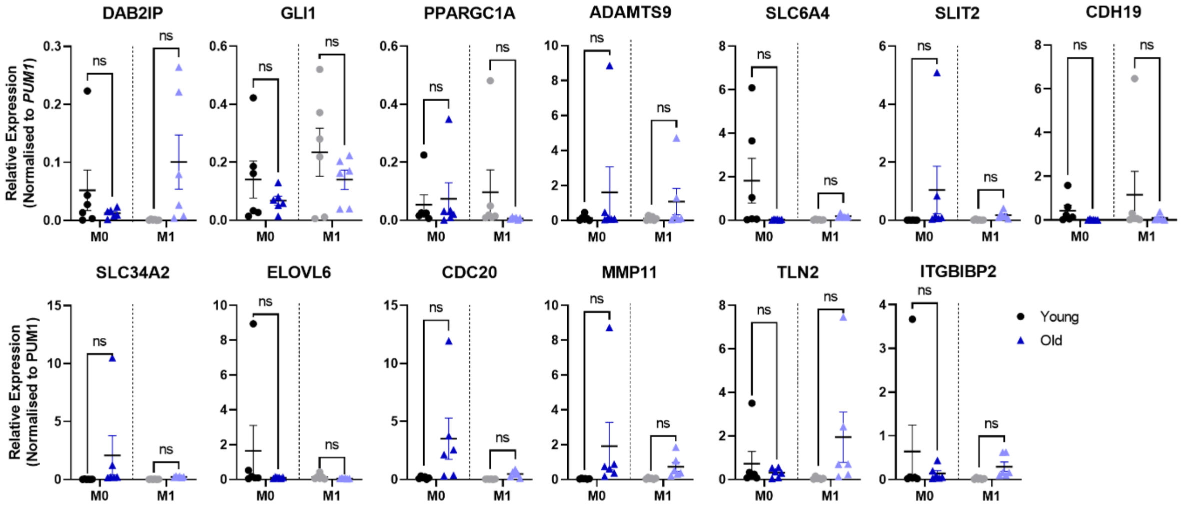

**Supplementary Fig 5 – Age-related changes in expression of genes of interest from RNA sequencing analysis of transcription factor knockdown**

Age-related changes in expression of selected genes in human monocyte derived macrophages isolated from young (22-25 years) and old (54-71 years) healthy donors that corresponded to genes dysregulated in transcription factor knockdown RNA sequencing comparisons. N = 6, unpaired T-test with Mann-Whitney test. *PUM1* expression was used as an internal control.

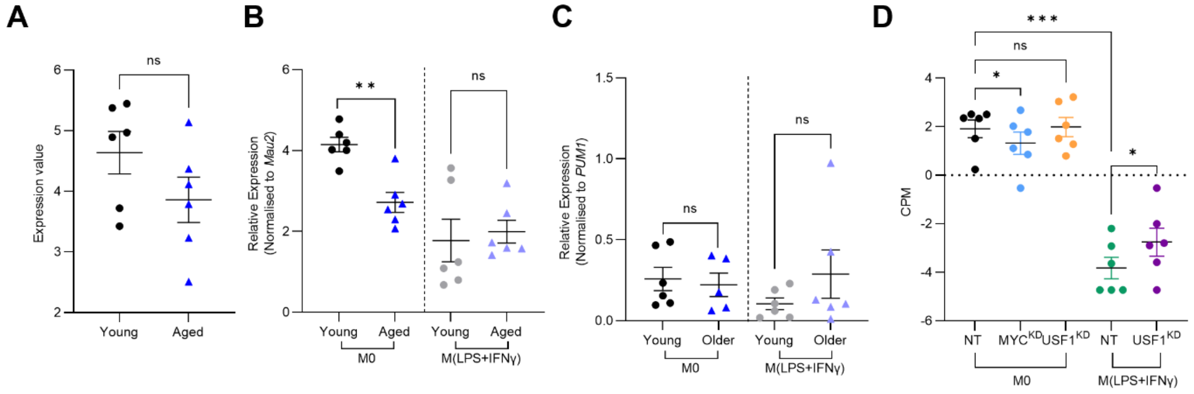


**Supplementary Fig 6 – *CCR2* expression in different murine and human macrophage populations**

A. Age-related changes in *CCR2* expression in murine alveolar macrophages via microarray analysis. N=6, Mann-Whitney test.
B. Age-related changes in *CCR2* expression in bone marrow-derived macrophages isolated from young (2-5 months) and aged (22-24 months) C57BL/6 mice by RT-qPCR. BMDMs were differentiated for 5 days with M-CSF and left unstimulated or further stimulated with LPS and IFNγ for 24 hours. N=6, Mann-Whitney test, ** P < 0.01. *Mau2* expression was used as an internal control.
C. Age-related changes in *CCR2* expression in human monocyte-derived macrophages isolated from young (22-25 years) and older (54-71 years) healthy donors by RT-qPCR. MDMs were differentiated for 7 days with M-CSF and left unstimulated or further stimulated with LPS and IFNγ for 24 hours. N=6, Mann-Whitney test. *PUM1* expression was used as an internal control.
D. Relative expression of *CCR2* after *MYC* or *USF1* knockdown in young human monocyte-derived macrophages via RNA sequencing analysis. MDMs were differentiated for 7 days with M-CSF and stimulated with siRNA for 72 hours. They were then left unstimulated or further stimulated with LPS and IFNγ for 24 hours. N=6, One-way ANOVA with Sidaks multiple comparison, * P < 0.05, *** P < 0.001.

**Supplementary Table 2 – Hallmarks associated with differentially expressed genes in si*MYC* $M^0$ vs $M^0$ control by GSEA analysis**

| Hallmark gene set | Size | ES | Nominal P value | Genes |
|---|---|---|---|---|
| Interferon alpha response | 56/95 | -0.55 | 0.000 | SP110, TDRD7, CMTR1, IFITM2, HLA-C, CCRL2, LGALS3BP, CASP8, SAMD9, IFI44, PARP14, B2M, DDX60, IL7, PARP12, PSMA3, MOV10, PSMB9, EIF2AK2, LAMP3, PARP9, EPSTI1, PSME2, BST2, MX1, PSMB8, CNP, HERC6, NUB1, IFITM3, PROCR, HELZ2, IFI35, TRIM21, CMPK2, IFIT3, RSAD2, ISG15, IFI44L, IFI27, IFIT2, LY6E, OASL, UBE2L6, PSME1, LAP3, TAP1, NMI, ISG20, USP18, TRAFD1, IFITM1, GMPR, IL15, CXCL11, GBP4 |
| MTORC1 signalling | 98/197 | -0.48 | 0.000 | HMGCR, SLC1A5, P4HA1, SLC7A11, PNP, PRDX1, ADD3, PSMA3, HSPD1, ATP5MC1, M6PR, ETF1, PPIA, PSAT1, CORO1A, HMGCS1, CDKN1A, RDH11, TM7SF2, TBK1, ARPC5L, ACTR3, PSMB5, EBP, DDX39A, G6PD, AURKA, SEC11A, ATP2A2, RPN1, HSPA9, SLC7A5, MAP2K3, ACLY, CCT6A, ACTR2, CALR, CFP, GLRX, GAPDH, RRM2, STARD4, PSMC2, GLA, PFKL, CACYBP, DHCR24, PPP1R15A, GBE1, CDC25A, SLC2A1, PPA1, PSME3, TOMM40, ENO1, RAB1A, HSPA5, DHCR7, FKBP2, LDLR, PSMC4, STIP1, HSP90B1, LTA4H, NFKBIB, TUBA4A, PSMD14, SQLE, TCEA1, MTHFD2, SERPINH1, IDI1, SLC9A3R1, RPA1, PSMA4, SC5D, ALDOA, ELOVL5, NUP205, PSPH, TUBG1, TES, NFIL3, SRD5A1, NUPR1, SYTL2, EIF2S2, ABCF2, HPRT1, SDF2L1, CYP51A1, MTHFD2L, HSPA4, ACSL3, STC1, POLR3G, ELOVL6, CTH |
| MYC target V1 | 100/195 | -0.45 | 0.000 | LDHA, MCM7, ODC1, RPL18, SF3B3, CCT2, SRPK1, NOLC1, EIF1AX, EIF4G2, RACK1, CDK4, RAD23B, SYNCRIP, PRDX3, EIF4E, UBE2E1, HNRNPA3, UBE2L3, HNRNPA2B1, HSPD1, PRPF31, PWP1, ETF1, PPIA, TOMM70, PSMA1, NDUFAB1, PRPS2, ILF2, YWHAQ, RPL22, CUL1, BUB3, RNPS1, AIMP2, RUVBL2, RPL14, SNRPA1, PSMA7, PSMD3, SRSF3, XRCC6, SNRPD2, CSTF2, PTGES3, TRIM28, HNRNPD, PCNA, PSMD8, SNRPB2, SET, SERBP1, MCM5, SSB, CBX3, PCBP1, HDGF, ERH, C1QBP, CCT5, PSMB3, SNRPD1, PSMD7, UBA2, NCBP2, CCNA2, CDK2, CCT3, PSMA6, HNRNPR, EIF2S1, GOT2, RANBP1, PSMC4, CYC1, COX5A, PSMD14, VDAC1, GLO1, KPNA2, MRPL23, CCT7, PSMD1, G3BP1, PSMA4, SNRPD3, HNRNPC, PRDX4, HSP90AB1, EXOSC7, PPM1G, EIF2S2, HPRT1, SSBP1, VBP1, TXNL4A, PSMB2, PSMA2, NCBP1 |
| Oxidative phosphorylation | 115/200 | -0.44 | 0.000 | COX7A2L, NQO2, MGST3, SLC25A5, ECH1, NDUFA3, VDAC3, DLAT, ATP5F1B, NDUFB7, UQCR10, ATP6V1D, SURF1, IMMT, ATP5MC3, COX8A, DECR1, SDHB, COX7A2, ATP5PD, COX6A1, NDUFB2, PDHB, SLC25A6, ACAA2, LDHA, PDHA1, COX5B, HTRA2, UQCRC1, ACAA1, SUPV3L1, NDUFC2, HCCS, SDHD, MRPS22, ACADM, MDH1, AIFM1, OGDH, GLUD1, PRDX3, ISCU, MPC1, ATP5MC1, UQCRB, TIMM50, COX7C, TOMM70, GRPEL1, NDUFS4, FXN, UQCRH, NDUFAB1, ATP5F1A, ATP1B1, LDHB, ETFB, HSPA9, ATP5F1D, MDH2, ATP6AP1, MRPS12, ATP5PF, OPA1, MRPS15, MTRR, ACAT1, SLC25A11, NDUFB4, ATP6V1G1, BAX, ATP6V0B, CYCS, GPX4, ACO2, COX4I1, ATP5F1E, MTX2, UQCRQ, FH, NDUFA9, UQCRFS1, MRPS11, TOMM22, ATP5F1C, HSD17B10, TIMM8B, GOT2, ATP6V1E1, CYC1, COX5A, VDAC2, NDUFB3, ATP6V0E1, NDUFS6, VDAC1, DLD, NDUFV2, CYB5R3, COX6C, ATP6V1F, COX17, TIMM17A, FDX1, COX11, TIMM10, CASP7, MRPL15, NNT, ATP6V0C, CYB5A, NDUFA8, COX6B1, ATP6V1H |
| TNF signaling via NFKB | 107/199 | -0.44 | 0.000 | RHOB, LAMB3, KLF6, TGIF1, CCRL2, JUNB, TNIP2, JAG1, CD80, DNAJB4, IL15RA, ICAM1, EIF1, SLC16A6, TLR2, PNRC1, NFKBIA, CXCL2, CXCL3, DUSP5, G0S2, KLF4, TRIP10, PLEK, YRDC, EGR1, B4GALT5, TRAF1, CCL4, BIRC2, TNF, NINJ1, BIRC3, IL6, GCH1, BTG1, GEM, ACKR3, IER5, CDKN1A, CLCF1, BCL2A1, FOSL1, TRIB1, TNFAIP6, PLAUR, DUSP4, IER2, IER3, IL23A, MAP2K3, TNFAIP3, PMEPA1, GFPT2, KYNU, NFE2L2, CXCL1, B4GALT1, KLF9, NR4A3, MAFF, DUSP1, DRAM1, ATF3, PHLDA1, NFKB1, CD83, HES1, CSF2, PANX1, PPP1R15A, TSC22D1, FOS, EHD1, RCAN1, CCL2, ID2, TNFRSF9, TANK, IFIT2, LDLR, CCL20, CEBPB, DDX58, TAP1, MSC, IL12B, JUN, NR4A1, PTX3, SPHK1, SPSB1, FOSB, IFNGR2, SLC2A6, KLF2, TNFSF9, TUBB2A, NFKBIE, NFIL3, LIF, PHLDA2, RELB, BTG3, SDC4, CXCL11, PLK2 |
| Interferon gamma response | 87/198 | -0.41 | 0.007 | B2M, DDX60, ZBP1, IL7, PNP, PARP12, ZNFX1, CASP3, PSMA3, FAS, IL6, GCH1, BTG1, SOCS1, FCGR1A, CDKN1A, IRF4, PSMB10, CFB, TNFAIP6, PSMB9, PTPN6, EIF2AK2, CD274, TNFAIP3, OAS2, SLAMF7, EPSTI1, PSME2, GBP6, BPGM, BST2, MX1, CD40, MT2A, PSMB8, MVP, CCL7, HERC6, IFITM3, NFKB1, HELZ2, MYD88, LYSMD2, IFI35, STAT1, ITGB7, TRIM21, CMPK2, IFIT3, RSAD2, PML, VCAM1, ISG15, IDO1, IFI44L, CCL2, SRI, OAS3, AUTS2, IFI27, NLRC5, IFIT2, LY6E, OASL, IFIT1, UBE2L6, PSME1, DDX58, LAP3, TAP1, NUP93, NMI, MTHFD2, ISG20, USP18, TOR1B, TRAFD1, GPR18, CASP7, IL15, ISOC1, ARL4A, PSMB2, PSMA2, CXCL11, GBP4 |
| Androgen response | 41/99 | -0.4 | 0.027 | XRCC5, HOMER2, B2M, ANKH, FKBP5, HMGCR, IQGAP2, HMGCS1, MERTK, XRCC6, UBE2J1, PMEPA1, B4GALT1, ELK4, DHCR24, TSC22D1, ALDH1A3, SLC38A2, SMS, ACTN1, DBI, ELL2, PTPN21, CENPN, IDI1, MYL12A, SRP19, STK39, ADRM1, TPD52, ELOVL5, CDC14B, VAPA, KLK2, NKX3-1, MAP7, NGLY1, ACSL3, APPBP2, SPDEF, BMPR1B |
| G2M checkpoint | 86/195 | 0.43 | 0.031 | TRAIP, CDC20, SMAD3, PURA, ARID4A, KNL1, KIF20B, BUB1, CENPA, EGF, EXO1, PDS5B, NDC80, NUMA1, POLA2, CCNB2, RPS6KA5, SS18, STMN1, KIF23, SMC4, YTHDC1, MAD2L1, CDC7, ATRX, PLK4, SRSF10, MAP3K20, ORC6, RBM14, NUSAP1, RBL1, STAG1, CKS1B, TNPO2, CCNF, EZH2, CCNT1, POLE, ODF2, CDC6, PTTG3P, RASAL2, XPO1, NOTCH2, LBR, RACGAP1, KIF4A, KIF15, PBK, HMMR, E2F2, MTF2, CENPF, MKI67, TACC3, MCM3, CUL5, DBF4, MCM6, KIF11, PLK1, SFPQ, WRN, CDKN1B, CDC25B, INCENP, MYBL2, NUP50, DKC1, KIF22, PAFAH1B1, ESPL1, LIG3, SMARCC1, KMT5A, TROAP, EWSR1, CENPE, SAP30, TRA2B, PRPF4B, RAD54L, ABL1, ILF3, PTTG1 |

### Supplementary Table 3 – Hallmarks associated with differentially expressed genes in si*USF1* $M^0$ vs $M^0$ control by GSEA analysis

| Hallmark gene set | Size | ES | Nominal P value | Genes |
|---|---|---|---|---|
| Interferon alpha response | 57/95 | -0.6 | 0.000 | CSF1, UBE2L6, IRF9, SELL, B2M, LAMP3, PARP9, LAP3, DDX60, ADAR, TRIM5, STAT2, PLSCR1, OAS1, PARP14, RTP4, SP110, DHX58, SAMD9L, CCRL2, GMPR, BST2, EIF2AK2, IFI44, GBP4, UBA7, IFITM2, EPSTI1, IFI30, IL7, PSMB9, LGALS3BP, TRIM25, TAP1, HERC6, MX1, IFI35, IFITM1, OASL, CXCL10, IFIT3, ISG15, IFIT2, IFITM3, LY6E, USP18, CMPK2, ISG20, TRAFD1, RSAD2, CD74, LPAR6, IFI44L, PARP12, IFI27, NCOA7, CXCL11 |
| Interferon gamma response | 89/198 | -0.4 | 0.019 | PML, RIPK1, SAMHD1, UBE2L6, NOD1, PTPN1, IRF9, B2M, APOL6, LAP3, XAF1, CD274, NFKB1, DDX60, ICAM1, ADAR, STAT2, CFB, PLSCR1, ARL4A, STAT1, PARP14, RTP4, SP110, DHX58, SAMD9L, IRF4, RNF213, METTL7B, KLRK1, BST2, IDO1, EIF2AK2, JAK2, MT2A, IFI44, CFH, GBP4, CSF2RB, TOR1B, GPR18, IFITM2, GBP6, EPSTI1, GZMA, DDX58, IFI30, OAS3, MX2, IL7, PSMB9, SLAMF7, LGALS3BP, HLA-DRB1, CMKLR1, TRIM25, OAS2, TAP1, IFNAR2, HERC6, MX1, IFI35, OASL, CXCL10, IFIT3, ISG15, TNFAIP2, IFIT2, IFITM3, LY6E, USP18, CIITA, HLA-DMA, IL18BP, CMPK2, HLA-DQA1, IFIT1, ISG20, TRAFD1, RSAD2, IRF5, ZBP1, CD74, IFI44L, BANK1, PARP12, PIM1, IFI27, CXCL11 |
| MYC targets | 100/195 | 0.53 | 0.000 | DEK, PGK1, ORC2, CDC20, LDHA, GSPT1, PA2G4, ACP1, HDDC2, CTPS1, HDAC2, MCM5, COPS5, NHP2, CDC45, BUB3, SYNCRIP, IFRD1, MRPL9, SMARCC1, SRSF3, RAN, FBL, EIF2S1, TRA2B, U2AF1, PSMA4, MYC, HDGF, EIF3J, CCNA2, PSMA2, TYMS, HNRNPA3, CCT4, HPRT1, PSMC4, PSMD3, AP3S1, PSMA6, YWHAE, USP1, XPO1, HNRNPR, PRPS2, SET, PWP1, PPIA, LSM2, NME1, GNL3, POLD2, SNRPD2, RFC4, RAD23B, VDAC1, MCM6, TARDBP, SNRPD1, TCP1, APEX1, ETF1, HNRNPA1, RPS6, CCT7, RPS3, EIF3B, HNRNPA2B1, SSBP1, EIF1AX, SF3A1, EIF4H, PRDX3, ILF2, PSMC6, NPM1, SRM, RSL1D1, SNRPG, SRSF1, KPNA2, XRCC6, SNRPB2, RUVBL2, HSPE1, PSMA1, STARD7, CCT2, AIMP2, CDK2, MRPS18B, DDX21, PSMD7, FAM120A, EIF4E, CBX3, SERBP1, HSPD1, IMPDH2, CCT3 |
| Glycolysis | 103/198 | 0.5 | 0.000 | PPFIA4, DEPDC1, SAP30, CENPA, PLOD2, EGFR, STC2, BIK, SPAG4, TFF3, FBP2, NT5E, PGK1, DCN, PGAM1, COL5A1, FUT8, PAXIP1, MIF, TPI1, PAM, EGLN3, EFNA3, KDELR3, LDHA, ERO1A, AK4, VLDLR, PFKP, CHST2, B4GALT7, ENO2, IRS2, VCAN, B3GNT3, RPE, AGL, PKM, HS2ST1, ENO1, SLC16A3, STC1, NDST3, ALDH7A1, PYGL, QSOX1, ANGPTL4, PLOD1, CASP6, LHPP, ANKZF1, GALK2, COPB2, PGM2, KIF2A, CHST12, DDIT4, ALDH9A1, PSMC4, ALDOA, FAM162A, ADORA2B, IGFBP3, LHX9, HMMR, XYLT2, PC, NOL3, CHST6, POLR3K, GLCE, GMPPA, KIF20A, PPIA, CHST4, GMPPB, GALK1, GFPT1, ME2, B4GALT4, ARTN, GALE, CLDN9, PFKFB1, ME1, TPST1, ALG1, VEGFA, CITED2, IER3, AGRN, P4HA1, P4HA2, GUSB, PGAM2, ARPP19, PHKA2, B4GALT2, NDUFV3, SLC37A4, MERTK, DLD, GYS1 |
| Hypoxia | 78/197 | 0.46 | 0.000 | PPARGC1A, PPFIA4, CP, SAP30, AKAP12, PKP1, PCK1, INHA, EGFR, STC2, DTNA, LOX, ALDOC, PGK1, GCK, GPI, DCN, PFKFB3, COL5A1, KDM3A, BCAN, PGM1, MIF, ETS1, TPI1, PAM, BNIP3L, EFNA3, KDELR3, NR3C1, LDHA, ENO3, BGN, ERO1A, AK4, VLDLR, PRDX5, TMEM45A, PFKP, CHST2, SLC2A1, ENO2, PPP1R3C, IRS2, NDST1, TPD52, PDK1, B4GALNT2, KIF5A, SLC2A5, WSB1, PRKCA, ENO1, XPNPEP1, GCNT2, STC1, MYH9, ANGPTL4, BCL2, CASP6, SLC25A1, ANKZF1, ACKR3, PGM2, SIAH2, CA12, DDIT4, ALDOA, FAM162A, ADORA2B, IGFBP3, GAPDH, STBD1, NEDD4L, CAVIN1, TPST2, GBE1, SERPINE1 |
| MTORC1 signaling | 94/197 | 0.44 | 0.003 | PHGDH, BUB1, PLOD2, PGK1, HMGCR, GPI, IGFBP5, FADS1, SORD, PGM1, PITPNB, LDLR, TPI1, UNG, EGLN3, LDHA, CCNG1, SRD5A1, ADIPOR2, ERO1A, AK4, VLDLR, ACACA, SLC2A1, CORO1A, UFM1, COPS5, PDK1, BCAT1, ENO1, IFRD1, PSMC2, GMPS, DHCR7, STC1, NAMPT, PSMA4, UCHL5, PSMG1, MLLT11, DDIT4, HPRT1, PSMC4, ALDOA, DHFR, GAPDH, CYB5B, ATP6V1D, CCT6A, TMEM97, XBP1, SLC1A4, GBE1, PSMB5, SSR1, ARPC5L, CCNF, EEF1E1, CACYBP, ACSL3, PPIA, SERPINH1, ACTR2, ITGB2, ELOVL5, STARD4, PNP, SERP1, SCD, ETF1, PNO1, SLC2A3, FADS2, CYP51A1, EDEM1, PFKL, SEC11A, PSMC6, ME1, FDXR, YKT6, PSMD12, P4HA1, ADD3, TES, ACTR3, ATP2A2, HSPE1, SLC37A4, TUBG1, CD9, TCEA1, TM7SF2, HSPD1 |
| Unfolded protein response | 57/110 | 0.44 | 0.020 | STC2, DKC1, KDELR3, ERO1A, DDX10, CKS1B, NHP2, EIF2AK3, CNOT2, POP4, DCP2, SDAD1, EIF2S1, TTC37, MTREX, DDIT4, CXXC1, XBP1, SLC1A4, SSR1, EEF2, ERN1, EXOC2, BAG3, FUS, GEMIN4, CHAC1, DNAJB9, CNOT6, DNAJC3, SPCS3, ALDH18A1, SERP1, EIF4EBP1, LSM4, DCTN1, EDEM1, SEC11A, EXOSC5, NPM1, KIF5B, CNOT4, VEGFA, TUBB2A, EIF4A2, BANF1, NABP1, CEBPB, SHC1, PREB, TATDN2, EIF4E, LSM1, SRPRB, ZBTB17, PAIP1, NOLC1 |
| Oxidative phosphorylatio n | 105/200 | 0.42 | 0.013 | ACADSB, GPI, SLC25A12, NDUFB1, SLC25A4, LDHA, ALDH6A1, COX17, TIMM50, NDUFB6, COX10, MRPL11, SUCLG1, PDHX, UQCRB, NDUFC1, ECHS1, DLST, NDUFA9, NDUFS1, GRPEL1, MTRF1, ECH1, ATP5MF, ATP5F1E, UQCR11, FXN, TOMM22, CS, COX7A2L, MRPS22, RETSAT, NDUFA1, ATP6V1D, HTRA2, ATP5PD, ATP5F1C, NDUFB8, ETFDH, NDUFA2, ISCA1, AFG3L2, ATP5ME, UQCRFS1, VDAC1, ALAS1, ATP5MC3, ISCU, VDAC2, ACAA1, GLUD1, MRPL35, NDUFB3, COX6B1, NDUFA4, NDUFC2, ATP5MG, MTRR, PRDX3, COX7A2, PDP1, BAX, ETFA, NDUFB2, POLR2F, BDH2, ATP5F1A, NDUFA6, PMPCA, SLC25A11, TIMM13, NDUFS3, SDHA, TIMM17A, FH, MTX2, ATP5PF, DLD, NDUFB5, MFN2, ETFB, SDHC, HCCS, NDUFA5, RHOT2, COX7C, TOMM70, IDH2, NDUFS6, NDUFS8, BCKDHA, ACADM, CYC1, MDH1, ATP5F1B, SDHD, SLC25A6, UQCRQ, LRPPRC, GOT2, IDH3A, MGST3, TIMM8B, OGDH, PDHB |

### Supplementary Table 4 – Hallmarks associated with differentially expressed genes in si*USF1* $M^{LPS+IFN\gamma}$ vs $M^{LPS+IFN\gamma}$ control by GSEA analysis

| Hallmark gene set | Size | ES | Nominal P value | Genes |
|---|---|---|---|---|
| Hypoxia | 91/197 | -0.52 | 0.000 | HEXA, PDK3, HAS1, VLDLR, VHL, SLC2A1, TPD52, CAVIN3, TPBG, CHST3, GPC3, TES, JUN, CA12, GBE1, SRPX, SIAH2, PLAUR, GPC4, PPFIA4, CDKN1C, SLC6A6, KDELR3, DPYSL4, PKLR, PGF, PDK1, CASP6, SDC3, IL6, CAVIN1, TGM2, KLHL24, PRKCA, SCARB1, TGFBI, SLC2A5, WSB1, ETS1, NDST1, MAP3K1, VEGFA, SLC2A3, HS3ST1, PHKG1, ERO1A, FBP1, ENO2, DDIT3, PGM1, ILVBL, STC2, FOSL2, GAPDHS, KLF7, ACKR3, ATP7A, EDN2, NFIL3, HK2, IRS2, FOXO3, LARGE1, LDHA, CITED2, ANKZF1, TPST2, BTG1, TKTL1, SDC2, LOX, KIF5A, CP, COL5A1, KDM3A, ERRFI1, P4HA1, PCK1, CDKN1B, MXI1, DCN, BNIP3L, EFNA3, CAV1, CXCR4, DDIT4, TMEM45A, PPARGC1A, CCNG2, STC1, BCAN |
| Angiogenesis | 17/36 | -0.63 | 0.012 | TIMP1, JAG2, FGFR1, LRPAP1, ITGAV, VEGFA, LPL, NRP1, SPP1, APP, CXCL6, VCAN, TNFRSF21, FSTL1, OLR1, PF4, STC1 |
| Heme metabolism | 93/194 | -0.48 | 0.000 | SLC30A1, KHNYN, MPP1, HDGF, CTNS, MINPP1, SIDT2, SLC22A4, PPOX, TYR, PSMD9, HEBP1, DCAF11, FBXO34, ADIPOR1, CCND3, SLC2A1, CAST, DMTN, SDCBP, SMOX, MFHAS1, RBM5, TMCC2, TNS1, NFE2L1, RHD, FOXJ2, UROD, EPB41, IGSF3, FN3K, SYNJ1, HTRA2, ADD2, PICALM, ARL2BP, HTATIP2, HMBS, EIF2AK1, TCEA1, ACKR1, RNF123, MYL4, VEZF1, CA2, EPOR, GAPVD1, LMO2, LRP10, CLCN3, PC, LAMP2, NEK7, BMP2K, CTSB, SLC7A11, GDE1, EZH1, DAAM1, ABCG2, FOXO3, ALDH1L1, TFDP2, ASNS, FECH, CCDC28A, AGPAT4, TFRC, RANBP10, XK, ARHGEF12, ELL2, TRIM58, TRAK2, ALAD, ACSL6, KAT2B, YPEL5, DCUN1D1, HBQ1, CROCCP2, ALDH6A1, NUDT4, ERMAP, MXI1, BNIP3L, GATA1, MOSPD1, TSPAN5, CAT, SLC6A9, MBOAT2 |
| Bile acid metabolism | 49/112 | -0.51 | 0.000 | HSD17B4, GNMT, DHCR24, LONP2, CYP46A1, CYP8B1, PIPOX, AQP9, PEX13, ABCA6, HACL1, PXMP2, ABCD1, ATXN1, TFCP2L1, HSD17B6, PNPLA8, PECR, FADS2, HSD17B11, FADS1, PHYH, ABCD2, ALDH8A1, ALDH1A1, ABCA8, BMP6, HSD3B1, RXRA, SLC27A2, IDH2, IDH1, ABCA1, NR3C2, SLC29A1, SLC27A5, HAO1, NUDT12, AMACR, CYP7A1, PEX11G, ABCD3, PEX1, SLC23A1, ALDH9A1, PEX7, CAT, PEX19, ABCA4 |
| Mitotic spindle | 198 | -0.45 | 0.000 | NCK2, SMC1A, CLIP1, KIF1B, LATS1, RANBP9, FSCN1, CKAP5, FGD6, TBCD, ARHGEF2, SUN2, SPTAN1, SSH2, ARAP3, TPX2, CCDC88A, SMC3, EPB41, CCNB2, PCNT, DST, WASF2, CLIP2, STK38L, NIN, ROCK1, TRIO, RACGAP1, PALLD, NF1, TUBGCP3, GSN, TSC1, NCK1, PLK1, RABGAP1, DYNLL2, LLGL1, CENPF, PPP4R2, ALS2, ARHGEF3, ARFGEF1, ARHGAP29, BIRC5, KIF4A, MARK4, HOOK3, RHOF, ITSN1, RASAL2, DOCK2, VCL, FGD4, SPTBN1, CDK1, TUBA4A, EZR, ACTN4, SAC3D1, TLK1, NEDD9, SYNPO, ARHGEF12, WASF1, SOS1, WASL, GEMIN4, NDC80, PKD2, CNTRL, EPB41L2, CDC42BPA, APC, CD2AP, AURKA, NEK2, NUSAP1, MID1IP1, ABL1, ECT2, SASS6, SHROOM2 |
| Apical junction | 81/199 | -0.44 | 0.000 | RRAS,MMP2,ADAM23,FSCN1,ADRA1B,SIRPA,MPZL2,GNAI2,MPZL1,CNN2,TJP1,BAIAP2,CRB3,HRAS,NF2,CDSN,SYK,ITGA3,CRAT,SGCE,NF1,CDH4,ITGA2,LIMA1,SDC3,TSC1,ITGA9,TGFBI,PTPRC,CD99,COL17A1,INPPL1,CX3CL1,CLDN7,ALOX15B,JUP,MSN,CD34,NECTIN3,ITGB1,RHOF,NLGN2,CD86,GRB7,VCL,YWHAH,PDZD3,ACTN4,ADAM9,JAM3,B4GALT1,PCDH1,COL16A1,DSC3,PIK3CB,WASL,CERCAM,AKT3,VCAN,SLC30A3,GTF2F1,IRS1,MAPK14,TMEM8B,EPB41L2,CLDN6,EXOC4,PTEN,FBN1,PECAM1,MMP9,MDK,NECTIN1,AMIGO1,ARHGEF6,CADM3,PARVA,CDH8,ACTA1,SHROOM2,VWF |
| MTORC1 signaling | 89/197 | -0.42 | 0.010 | TXNRD1, SKAP2, LGMN, ALDOA, MLLT11, ME1, ACTR2, STIP1, PSME3, IGFBP5, CACYBP, SC5D, VLDLR, SLC7A5, SLC2A1, DHCR24, SCD, TES, FGL2, ADIPOR2, GBE1, CDC25A, SLC1A5, CFP, MTHFD2, STARD4, CYP51A1, EIF2S2, SQLE, EBP, HMGCR, SLC6A6, RRM2, BCAT1, SSR1, ITGB2, HMBS, HMGCS1, CANX, PDK1, UNG, TCEA1, LDLR, CORO1A, PLOD2, ACLY, PLK1, DHFR, CTH, IFRD1, SLC2A3, CALR, RPA1, ERO1A, DDIT3, SLC7A11, PGM1, SLC9A3R1, LTA4H, FADS2, PHGDH, ATP2A2, SLC1A4, NAMPT, RDH11, FADS1, NFIL3, HK2, LDHA, TUBA4A, ASNS, CD9, TFRC, SORD, TRIB3, IDH1, PSPH, SHMT2, PSAT1, UCHL5, DHCR7, GOT1, AURKA, P4HA1, ELOVL6, CXCR4, DDIT4, ADD3, STC1 |
| MYC targets V2 | 35/57 | 0.61 | 0.000 | PPAN, NOP16, AIMP2, PES1, NOP2, RRP9, GNL3, SUPV3L1, HSPE1, RABEPK, MRTO4, IMP4, IPO4, PUS1, NPM1, NOLC1, MPHOSPH10, NOC4L, DUSP2, PPRC1, WDR43, PA2G4, WDR74, MYBBP1A, SRM, UTP20, RCL1, TBRG4, TCOF1, MCM4, HSPD1, NOP56, NDUFAF4, NIP7, CBX3 |
| MYC targets V1 | 111/195 | 0.50 | 0.000 | CDC20, PSMC6, SNRPA, NME1, NOP16, AIMP2, MRPL9, RRP9, GNL3, HSPE1, NHP2, ODC1, RPS10, EIF3B, SNRPD2, MRPL23, ABCE1, LSM7, DUT, TCP1, HDDC2, SNRPG, PSMB3, IMPDH2, FBL, C1QBP, CCT2, SNRPD1, NPM1, RANBP1, EEF1B2, RAN, SRSF7, ORC2, NOLC1, PSMA2, EIF3D, PRPS2, NDUFAB1, CCT3, PA2G4, CDC45, SRSF2, PTGES3, SRM, DHX15, TRA2B, RFC4, COX5A, PCNA, CSTF2, CTPS1, SNRPA1, TUFM, TOMM70, PSMD3, MCM4, HDAC2, RPS5, HSPD1, CUL1, CYC1, ILF2, SNRPB2, NOP56, TRIM28, PHB2, EIF4A1, PSMA1, ERH, RPS2, VBP1, APEX1, PSMA4, CCT5, CCT7, HNRNPA2B1, SNRPD3, CBX3, HNRNPC, CAD, HSP90AB1, RNPS1, EIF3J, TXNL4A, XRCC6, HNRNPR, RUVBL2, MYC, PPIA, HNRNPA1, PSMD14, RPL14, POLD2, LSM2, AP3S1, POLE3, DDX21, RPL6, SSB, RPL18, VDAC1, STARD7, PSMA7, PSMA6, SF3B3, UBE2E1, PSMD1, PSMD8, RPS6, SRSF3 |

**Supplementary Table 5 – Enriched biological processes associated with differentially expressed genes between young (2-4 months) and old (22-24 months) alveolar macrophages isolated from C57BL/6J mice**

| Biological process | Gene count | Adjusted P value | Genes |
|---|---|---|---|
| Chromosome segregation | 52/324 | 2.35E-10 | Cpeb1; Ska1; Cdc20; Incenp; Anapc1; Dscc1; Birc5; Cenpq; Ube2c; Brca1; Ncapd2; Ect2; Ccnb1; Nusap1; Cenpt; Hjurp; Bub1b; Atrx; Plk1; Cenpn; Cdca8; Mis18a; Cenpk; Nipbl; Kif23; Nuf2; Uvrag; Top2a; Fbxo5; Usp9x; Prc1; Nsl1; Nup37; Chmp1a; Sirt7; Cenpf; Mad2l1; Bub1; Spag5; Fmn2; Chmp6; Aurkb; Dsn1; Ska3; Mms19; Cep192; Cenpw; Ncaph; Ncapg; Knstrn; Kif4; Oip5 |
| Mitotic cell cycle phase transition | 56/376 | 3.17E-10 | Cdkn2a; Ube2e2; Ccnb2; Pdpn; Cdkn2b; Anapc1; Zfp36l1; Tcf19; Birc5; Cks1b; Tm4sf5; Plk2; Ube2c; Brca1; Ccnb1; Cdkn2c; Tjp3; Hinfp; Bub1b; Akt1; Cdc25c; Cdk4; Psme3; Pole; Plk1; Plrg1; Fhl1; Ptpn6; Cdk1; Mtbp; Pten; Ercc3; Chek1; Usp47; Fbxo5; FoxM$^{LPS+IFN\gamma}$; Mepce; Brsk1; Sirt7; Cenpf; Mad2l1; Rfwd3; Bub1; Ticrr; Iqgap3; Aven; Cacul1; Dtl; Prmt2; Cdk5rap3; Aurkb; Cep192; Anxa1; Rdx; Cdk2; Cdc25a |
| Regulation of mitotic cell cycle phase transition | 44/265 | 2.16E-09 | Cdkn2a; Ube2e2; Pdpn; Cdkn2b; Zfp36l1; Birc5; Tm4sf5; Ube2c; Brca1; Ccnb1; Tjp3; Bub1b; Akt1; Cdc25c; Cdk4; Psme3; Plk1; Plrg1; Fhl1; Ptpn6; Cdk1; Mtbp; Pten; Ercc3; Chek1; Usp47; Fbxo5; Mepce; Brsk1; Cenpf; Mad2l1; Rfwd3; Bub1; Ticrr; Aven; Dtl; Prmt2; Cdk5rap3; Aurkb; Cep192; Anxa1; Rdx; Cdk2; Cdc25a |
| Mitotic sister chromatid segregation | 32/151 | 2.39E-09 | Cdc20; Incenp; Anapc1; Dscc1; Birc5; Ube2c; Ncapd2; Ccnb1; Nusap1; Bub1b; Atrx; Plk1; Cdca8; Cenpk; Nipbl; Kif23; Nuf2; Fbxo5; Prc1; Nsl1; Chmp1a; Mad2l1; Bub1; Spag5; Chmp6; Aurkb; Dsn1; Cep192; Ncaph; Ncapg; Knstrn; Kif4 |
| Cell cycle phase transition | 57/415 | 2.39E-09 | Cdkn2a; Ube2e2; Ccnb2; Pdpn; Cdkn2b; Anapc1; Zfp36l1; Tcf19; Birc5; Cks1b; Tm4sf5; Plk2; Ube2c; Brca1; Ccnb1; Cdkn2c; Tjp3; Hinfp; Bub1b; Akt1; Cdc25c; Cdk4; Psme3; Pole; Plk1; Plrg1; Fhl1; Ptpn6; Cdk1; Mtbp; Pten; Ercc3; Chek1; Usp47; Fbxo5; FoxM$^{LPS+IFN\gamma}$; Mepce; Brsk1; Sirt7; Cenpf; Mad2l1; Rfwd3; Bub1; Paf1; Ticrr; Iqgap3; Aven; Cacul1; Dtl; Prmt2; Cdk5rap3; Aurkb; Cep192; Anxa1; Rdx; Cdk2; Cdc25a |
| Mitotic nuclear division | 43/268 | 6.31E-09 | Igf1; Cdc20; Incenp; Pdgfb; Anapc1; Dscc1; Birc5; Ube2c; Ncapd2; Ccnb1; Nusap1; Il1b; Bub1b; Atrx; Plk1; Cdca8; Cenpk; Fbxw5; Nipbl; Mtbp; Kif11; Chek1; Kif23; Nuf2; Fbxo5; Prc1; Nsl1; Aaas; Chmp1a; Sirt7; Mad2l1; Bub1; Ereg; Spag5; Chmp6; Aurkb; Dsn1; Cep192; Ncaph; Ncapg; Knstrn; Kif4; Nme6 |
| Regulation of cell cycle phase transition | 45/297 | 1.31E-08 | Cdkn2a; Ube2e2; Pdpn; Cdkn2b; Zfp36l1; Birc5; Tm4sf5; Ube2c; Brca1; Ccnb1; Tjp3; Bub1b; Akt1; Cdc25c; Cdk4; Psme3; Plk1; Plrg1; Fhl1; Ptpn6; Cdk1; Mtbp; Pten; Ercc3; Chek1; Usp47; Fbxo5; Mepce; Brsk1; Cenpf; Mad2l1; Rfwd3; Bub1; Paf1; Ticrr; Aven; Dtl; Prmt2; Cdk5rap3; Aurkb; Cep192; Anxa1; Rdx; Cdk2; Cdc25a |
| Sister chromatid segregation | 33/181 | 4.14-08 | Cdc20; Incenp; Anapc1; Dscc1; Birc5; Ube2c; Ncapd2; Ccnb1; Nusap1; Bub1b; Atrx; Plk1; Cdca8; Cenpk; Nipbl; Kif23; Nuf2; Top2a; Fbxo5; Prc1; Nsl1; Chmp1a; Mad2l1; Bub1; Spag5; Chmp6; Aurkb; Dsn1; Cep192; Ncaph; Ncapg; Knstrn; Kif4 |
| Negative regulation of cell cycle | 56/451 | 1.12E-07 | Cdkn2a; Apbb2; Cdkn3; Cdkn2b; Pmp22; Zfp36l1; Birc5; H2-M3; Plk2; Sgsm3; Brca1; Myc; Ppp1r10; Ccnb1; Cdkn2c; Btg1; Kntc1; Hinfp; Dna2; Bub1b; Atrx; Cdk4; Cdk5rap1; Plk1; Nsun2; Fhl1; Cdk1; Cdk9; Mtbp; Pten; Trim35; Chek1; Wee1; Mdm4; Usp47; Top2a; Fbxo5; FoxM$^{LPS+IFN\gamma}$; Chmp1a; Gadd45a; Brsk1; Mad2l1; Rfwd3; Bub1; Map2k1; Ticrr; Nabp2; Aven; Dtl; Prmt2; Cdk5rap3; Aurkb; Gas2l1; Cep192; Trrap; Nme6 |
| Mitotic cytokinesis | 19/69 | 2.83E-07 | Kif20a; Snx18; Cenpa; Stmn1; Incenp; Birc5; Ckap2; Ect2; Nusap1; Plk1; Sptbn1; Unc119; Anln; Kif23; Prc1; Chmp1a; Chmp6; Kif4; Usp8 |
| Nuclear chromosome segregation | 39/262 | 3.15E-07 | Cpeb1; Cdc20; Incenp; Anapc1; Dscc1; Birc5; Cenpq; Ube2c; Ncapd2; Ect2; Ccnb1; Nusap1; Bub1b; Atrx; Plk1; Cdca8; Cenpk; Nipbl; Kif23; Nuf2; Top2a; Fbxo5; Prc1; Nsl1; Chmp1a; Sirt7; Cenpf; Mad2l1; Bub1; Spag5; Fmn2; Chmp6; Aurkb; Dsn1; Cep192; Ncaph; Ncapg; Knstrn; Kif4 |
| Negative regulation of mitotic cell cycle | 36/236 | 6.65E-07 | Cdkn2b; Zfp36l1; Birc5; Plk2; Brca1; Ppp1r10; Ccnb1; Btg1; Kntc1; Bub1b; Atrx; Plk1; Fhl1; Cdk1; Mtbp; Pten; Trim35; Chek1; Wee1; Usp47; Top2a; Fbxo5; Gadd45a; Brsk1; Mad2l1; Rfwd3; Bub1; Ticrr; Nabp2; Aven; Prmt2; Cdk5rap3; Aurkb; Cep192; Trrap; Nme6 |
| Cytoskeleton-dependent cytokinesis | 21/94 | 1.76E-06 | Kif20a; Snx18; Cenpa; Stmn1; Incenp; Birc5; Ckap2; Ect2; Nusap1; Plk1; Sptbn1; Unc119; Anln; Kif23; Prc1; Chmp1a; Fmn2; Chmp6; Aurkb; Kif4; Usp8 |
| Regulation of cell-cell adhesion | 51/427 | 1.78E-06 | Pla2g2d; Cdkn2a; Cd74; H2-Aa; Pdpn; Igf1; CeacaM$^{LPS+IFN\gamma}$; H2-Ab1; Il6st; Itga6; Nfkbid; L1cam; H2-M3; Gm5150; Trpv4; Jak3; Malt1; Btn1a1; Il1b; Akt1; Icosl; Xbp1; Zfp608; Ptpn6; Dmtn; Tnfrsf21; Sh2b3; Cela2a; Cblb; Irak1; Rap1gap; Traf6; Ass1; Fut4; Bmi1; Fermt3; Ptafr; Erbb2; Nlrp3; Itch; Nr4a3; Map2k1; Anxa1; Cyld; Akna; Rdx; VcaM$^{LPS+IFN\gamma}$; IcaM$^{LPS+IFN\gamma}$; Specc1l; Ccl5; Nck2 |
| DNA repair | 55/481 | 1.99E-06 | Zranb3; Mcm3; Nudt16l1; Terf2ip; Brca1; Recql; Ap5s1; Mcm5; Mms22l; Hinfp; Dna2; Actl6a; Mcm2; Parp2; Parp1; Atrx; Trim28; Fbxo6; Pole; Gins2; Huwe1; Cdk7; Pole2; Cdk9; Rad51; Nipbl; Ercc3; Chek1; Dmap1; Morf4l2; Usp47; Eid3; Uvrag; FoxM$^{LPS+IFN\gamma}$ Ube2v1; Smug1; Exo1; Rfc3; Sirt7; Rad51ap1; Nfrkb; Rfwd3; Fmn2; Fancg; Ticrr; Gtf2h2; Nabp2; Dtl; Neil3; Wdr48; Mms19; Bod1l; Tdp2; Cdk2; Trrap |

**Supplementary Table 6 – Hallmarks associated with differentially expressed genes in old vs young C57BL/6J mouse alveolar macrophages**

| Hallmark gene sets | Size | ES | Nominal P value | Genes |
|---|---|---|---|---|
| MYC targets V1 | 78/126 | -0.49 | 0.049 | ODC1, TUFM, KARS1, CLNS1A, SNRPD1, CCT7, NME1, XRCC6, SNRPG, EIF4H, C1QBP, TFDP1, HDAC2, PHB1, SRSF7, PRDX3, CTPS1, SNRPD3, EIF2S1, PSMA6, RAN, KPNA2, DDX21, H2AZ1, GOT2, ORC2, HNRNPC, EIF4E, SERBP1, PCNA, CCT2, CYC1, PA2G4, PHB2, MCM7, RRP9, UBE2E1, MCM4, CSTF2, TXNL4A, MCM6, CNBP, TYMS, USP1, SLC25A3, RUVBL2, PSMC6, RACK1, PSMD1, GNL3, SNRPA1, RNPS1, NDUFAB1, SRSF3, TCP1, IARS1, NOP16, PRPS2, VBP1, KPNB1, IMPDH2, CCT3, CDK4, NCBP1, DUT, PPM1G, MYC, TOMM70, EXOSC7, PRPF31, LSM2, MRPS18B, MRPL9, CDK2, CDC45, TARDBP, AIMP2, MAD2L1 |
| G2m checkpoint | 51/116 | -0.57 | 0.022 | MKI67, HIRA, LMNB1, ARID4A, ORC6, UCK2, RACGAP1, E2F4, JPT1, TACC3, STMN1, AURKA, TROAP, KIF22, KPNB1, HMMR, HUS1, HMGB3, RASAL2, SMC2, CDK4, RBL1, TENT4A, MEIS1, PLK1, ODF2, POLE, CHAF1A, NEK2, MYC, NDC80, STIL, BRCA2, BIRC5, UBE2C, FBXO5, PLK4, EXO1, BUB1, NSD2, MCM3, CDC45, CDC6, CENPA, KIF23, TOP2A, NUSAP1, MAD2L1, KIF11, CDKN3, PBK |
| E2F targets | 76/130 | -0.57 | 0.014 | DNMT1, POLE4, UBR7, STAG1, MCM7, TUBG1, PRIM2, MCM4, CIT, DONSON, MCM6, USP1, MYBL2, CENPE, MKI67, EXOSC8, TK1, LMNB1, RAD50, ORC6, TIPIN, NUDT21, ZW10, GINS4, RPA3, TRIP13, RACGAP1, PMS2, JPT1, TACC3, STMN1, CHEK2, AURKA, E2F8, KIF22, PPM1D, TIMELESS, HMMR, SNRPB, HUS1, CNOT9, UBE2T, CENPM, HMGB3, CDK4, DUT, MELK, PLK1, TCF19, CDCA3, TBRG4, POLE, NBN, CSE1L, MYC, CCP110, PRKDC, BRCA2, BUB1B, BIRC5, ANP32E, PLK4, RAD1, MCM3, PSIP1, SPAG5, SHMT1, TOP2A, MAD2L1, PSMC3IP, SLBP, WEE1, BRCA1, DSCC1, CDKN3, CDCA8 |
| Spermatogenesis | 25/51 | -0.46 | 0.020 | ZC3H14, STRBP, TCP11, ZPBP, PSMG1, TSN, GMCL1, PIAS2, SCG5, RAD17, PHF7, TNP2, AURKA, COIL, PRKAR2A, PCSK1N, POMC, NEK2, MTOR, RPL39L, MLF1, BUB1, NCAPH, PARP2, CDKN3 |
| Epithelial mesenchymal transition | 31/94 | 0.53 | 0.030 | COL4A2, COL4A1, DPYSL3, SLC6A8, FSTL1, ECM1, GADD45A, GPC1, ANPEP, MYL9, ITGB3, DST, FGF2, AREG, CXCL12, VEGFA, ID2, MMP14, FAP, LAMC2, SDC4, SERPINE1, FLNA, SPP1, IGFBP4, SNTB1, ACTA2, CADM1, PVR, SAT1, LUM |
| Allograft rejection | 30/110 | 0.47 | 0.035 | CDKN2A, CXCL9, CSF1, EREG, PRKCB, CCL7, PF4, HLA-DQA1, ICOSLG, ITGAL, NLRP3, CXCR3, EIF4G3, HLA-DOA, IL13, ETS1, CD74, STAB1, GBP2, AKT1, HLA-DMB, HLA-G, CXCL13, IL10, CARTPT, CCL2, CCL4, SOCS1, FLNA, CD3E |

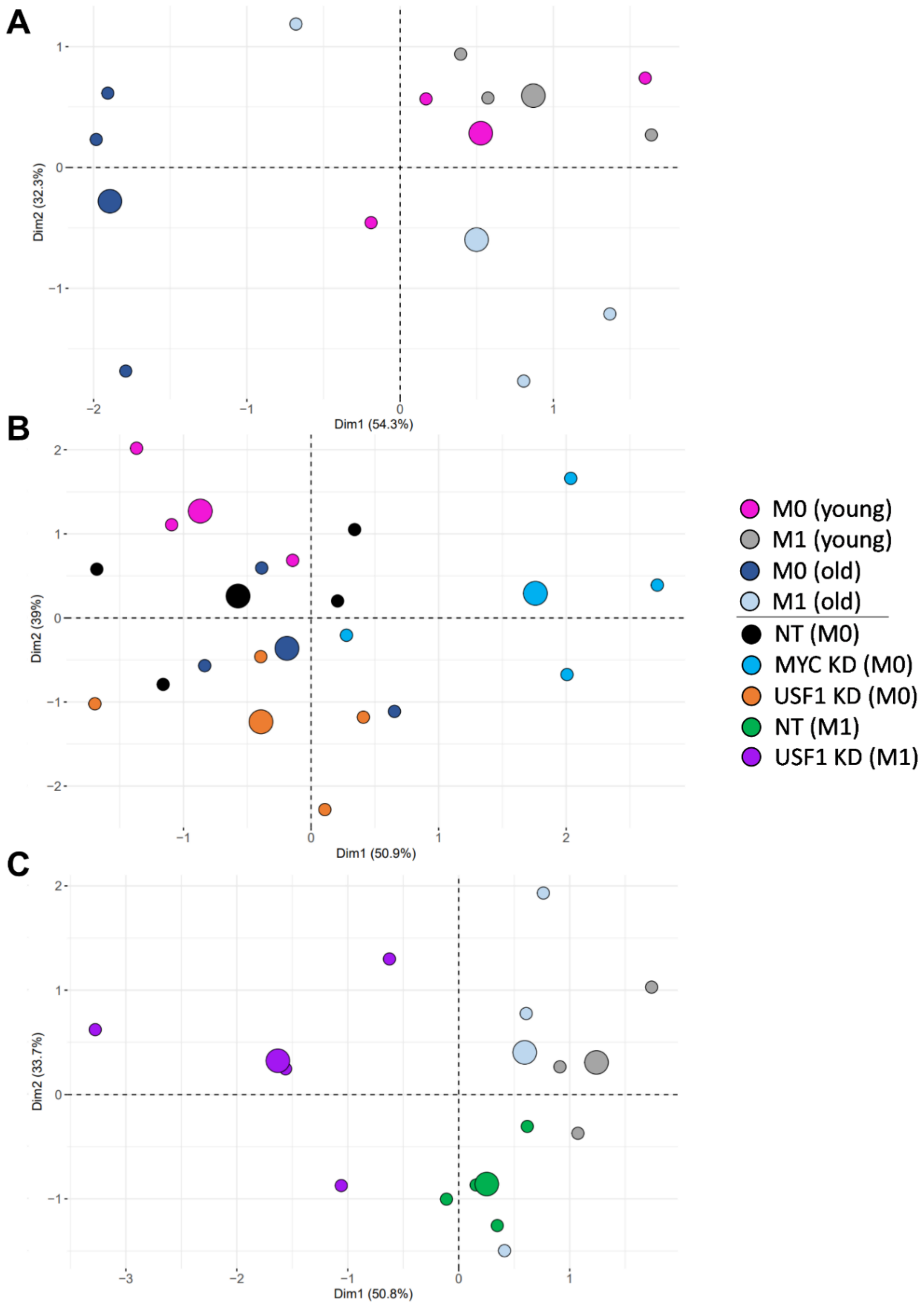


**Supplementary Fig 7 – PCA plots for cell morphology and actin staining**

Principal component analysis clustering samples by cell size, circularity and mean fluorescence intensity using prcomp in R.

A. Young and old samples.

B. All $M^0$ samples.

C. All $M^{LPS+IFN\gamma}$ samples.

**Supplementary Table 7 – Oligonucleotide sequences for RT-qPCR of human monocyte-derived macrophages**

| Target gene | Forward primer (5'-to-3') | Reverse primer (5'-to-3') |
|---|---|---|
| *PUM1* | GCATTTGGACAAGGTCTGGCAG | GCTACAAGTCGAACAGGAGCTC |
| *MYC* | AGAGTTTCATCTGCGACCCG | GAAGCCGCTCCACATACAGT |
| *USF1* | GCTCTATGGAGAGCACCAAGTC | AGACAAGCGGTGGTTACTCTGC |
| *NFYB* | GGAATTGGTGGAGCAGTCACAG | CCGTCTGTGGTTATTAAGCCAGC |
| *SRF* | TCACCTACCAGGTGTCGGAGTC | GTGCTGTTTGGATGGTGGAGGT |
| *FOXM1* | TCTGCCAATGGCAAGGTCTCCT | CTGGATTCGGTCGTTTCTGCTG |
| *NFIC* | TGGCGGCGATTACTACACTTCG | GGCTGTTGAATGGTGACTTGTCC |
| *ATP8B1* | CTTCTTGCTCGCAGTTTGCCAC | GCCAAAGTTCCTGGCAGCGTTT |
| *SFRP4* | CTATGACCGTGGCGTGTGCATT | GCTTAGGCGTTTACAGTCAACATC |
| *GDF1* | GTCACCCTGCAACCGTGCCAC | AGGTCGAAGACGACTGTCCACT |
| *STAB2* | ACTGGCTCCTTACCAAACCTGC | GAGCAAACACTGTGTAGGCATCG |
| *MMP8* | CAACCTACTGGACCAAGCACAC | TGTAGCTGAGGATGCCTTCTCC |
| *PLOD2* | GACAGCGTTCTCTTCGTCCTCA | CTCCAGCCTTTTCGTGGTGACT |
| *ITGB6* | TCTCCTGCGTGAGACACAAAGG | GAGCACTCCATCTTCAGAGACG |
| *ITGA2B* | CTGTCCAGCTACTGGTGCAAGA | ATGTTGTGCCCAGTGGCTCCAA |
| *CDH8* | AACGCTGGCAACACCACTTGAC | GCGTTGTCATTGACATCCAGCAC |
| *ADAMTS9* | CCATTCAGAGGTGCAGTGAGTTC | ACCAGACCTGGCGGTGCTTATG |
| *AJUBA* | AGCCACCAGGTCCTTTCGTTCC | GGCATTGCTCTGCCCATAGATG |
| *CDH19* | ATTGGTCAGCCAGGAGCGTTGT | GCAGATTCAGAGACAGTCAAGCG |
| *ANGPTL3* | CCTGAAACTCCAGAACACCCAG | TTCCACGGTCTGGAGAAGGTCT |
| *PCDHGB4* | CAACACGGACTGGCGTTTCTCT | GATCATGGCTTGCAGCATCTCTG |
| *DAB2IP* | TCATCGCCAAGGTCACCCAGAA | CGCTGCATGTTGGTCCACTCAT |
| *GLI1* | AGCCTTCAGCAATGCCAGTGAC | GTCAGGACCATGCACTGTCTTG |
| *PPARGC1A* | CCAAAGGATGCGCTCTCGTTCA | CGGTGTCTGTAGTGGCTTGACT |
| *SLC6A4* | TCACAGTGCTCGGTTACATGGC | GAAAGTGGACGCTGGCATGTTG |
| *SLIT2* | CAGAGCTTCAGCAACATGACCC | GAAAGCACCTTCAGGCACAACAG |
| *SLC34A2* | CGTGTGTGCATGGGTCAAAG | CAATCTTGCTGCACGGCTAC |
| *ELOVL6* | CCATCCAATGGATGCAGGAAAAC | CCAGAGCACTAATGGCTTCCTC |
| *CDC20* | CGGAAGACCTGCCGTTACATTC | CAGAGCTTGCACTCCACAGGTA |
| *MMP11* | GAGAAGACGGACCTCACCTACA | CTCAGTAAAGGTGAGTGGCGTC |
| *MMP13* | GCACTTCCCACAGTGCCTAT | AGTTCTTCCCTTGATGGCCG |
| *TLN2* | CAAGGAAGTCGCCAACAGCACT | TTGAGGCGAACGCTGTCAGGTT |
| *ITGB1BP2* | GACCACACTGTGCTGAGAAGCT | AGCAGCTTCAGAGGCAACTCTG |
| *WNT11* | CTGTGAAGGACTCGGAACTCGT | AGCTGTCGCTTCCGTTGGATGT |
| *GPR32* | CTGGGGCCCTTAGCAATCAT | AGATGGACCAACAGCACCAC |
| *CCR2* | GGGATGACTCACTGCTGCAT | TGCTTTCGGAAGAACACCGA |

**Supplementary Table 8 – Oligonucleotide sequences for RT-qPCR of C57BL6 mouse bone marrow-derived macrophages**

| Target gene | Forward primer (5'-to-3') | Reverse primer (5'-to-3') |
|---|---|---|
| *Mau2* | TGGTTACCTGGAGAAGGCACAG | ATGCTCCAGCAGGATCACTTGG |
| *Myc* | TTGAAGGCTGGATTTCCTTTGGGC | TCGTCGCAGATGAAATAGGGCTGT |
| *Usf1* | CGTCTTCCGAACTGAGAATGGG | CTGGGTCATAGACTGAGTGGCA |
| *Nfyb* | ACCAAACAGCCGATTGGAGA | CTAGCTGGGAGGCATCTGTG |
| *Srf* | CACCTACCAGGTGTCGGAAT | GTCTGGATTGTGGAGGTGGT |
| *Foxm1* | GTCTCCTTCTGGACCATTCACC | GCTCAGGATTGGGTCGTTTCTG |
| *Nfic* | TGACTCAGTAAGTTCGGCGG | GTTGAACCAGGTGTAGGCGA |
| *Ccr2* | GCTGTGTTTGCCTCTCTACCAG | CAAGTAGAGGCAGGATCAGGCT |
